# Pan-cancer Analysis of Homologous Recombination Deficiency in Cell Lines

**DOI:** 10.1101/2024.05.16.594524

**Authors:** Anne E. Dodson, Sol Shenker, Pamela Sullivan, Chris Middleton, Michael McGuire, Edmond Chipumuro, Yuji Mishina, Erica Tobin, Louise Cadzow, Andrew A. Wylie, Dipen Sangurdekar

## Abstract

Homologous Recombination Deficiency (HRD) drives genomic instability in multiple cancer types and renders tumors vulnerable to certain DNA damaging agents such as PARP inhibitors. Thus, HRD is emerging as an attractive biomarker in oncology. A variety of in silico methods are available for predicting HRD; however, few of these methods have been applied to cell lines in a comprehensive manner. Here we utilized two of these methods, “CHORD” and “HRDsum” scores, to predict HRD for 1,332 cancer cell lines and 84 non-cancerous cell lines. Cell lines with biallelic mutations in *BRCA1* or *BRCA2*, which encode key components of the homologous recombination pathway, showed the strongest HRD predictions, validating the two methods in cell lines. A small subset of *BRCA1/2*-wildtype cell lines were also classified as HRD, several of which showed evidence of epigenetic *BRCA1* silencing. Similar to HRD in patient samples, HRD in cell lines was associated with p53 loss, was mutually exclusive with microsatellite instability and occurred most frequently in breast and ovarian cancer types. In addition to validating previously identified associations with HRD, we leveraged cell line-specific datasets to gain new insights into HRD and its relation to various genetic dependency and drug sensitivity profiles. We found that in cell lines, HRD was associated with sensitivity to PARP inhibition in breast cancer, but not at a pan-cancer level. By generating large-scale, pan-cancer datasets on HRD predictions in cell lines, we aim to facilitate efforts to improve our understanding of HRD and its utility as a biomarker.

**SIGNIFICANCE STATEMENT:** Homologous Recombination Deficiency (HRD) is common in cancer and can be exploited therapeutically, as it sensitizes cells to DNA damaging agents. Here we scored over 1,300 cancer cell lines for HRD using two different bioinformatic approaches, thereby enabling large-scale analyses that provide insight into the etiology and features of HRD.

## INTRODUCTION

Homologous recombination (HR) is a high-fidelity mechanism for the repair of double-strand breaks. In the absence of HR, cells rely on more error-prone pathways such as polymerase theta-mediated end joining (TMEJ) to repair double-strand breaks and thereby accumulate specific types of mutations over time (Koh et al., 2021). The genomic instability that develops in response to HR deficiency (HRD) can, in turn, drive tumorigenesis. HRD is present in up to half of all high-grade serous ovarian cancers, at least a quarter of primary breast cancers, and a subset of pancreatic and prostate cancers (Konstantinopoulos et al., 2015; Leibowitz et al., 2022; L. Nguyen et al., 2020). As a source of genomic instability, HRD also sensitizes cells to certain types of DNA damage. Inhibition of poly(ADP-ribose) polymerase (PARP) leads to an accumulation of DNA lesions that are insufficiently repaired in the absence of BRCA1 or BRCA2, key components of the HR pathway (Bryant et al., 2005; Farmer et al., 2005). As a result, PARP inhibition leads to cell death in *BRCA1/2*-deficient tumors.

Due to its prevalence in cancer, HRD has garnered much interest from a clinical perspective, and several methods have been developed to detect HRD (Ngoi & Tan, 2021). Traditionally, tests have focused on germline and somatic mutations in *BRCA1* and *BRCA2*, the most common causes of HRD. Gene panels that include additional HR genes such as *BRIP1*, *PALB2*, and *RAD51C* have also been developed to capture potential causes of HRD that extend beyond *BRCA1/2* mutations (Yamamoto & Hirasawa, 2021). Although genetic testing is well-established, this approach has a few caveats. First, variants of unknown significance can be difficult to interpret. Second, genetic testing does not capture epigenetic alterations such as *BRCA1* promoter methylation, a mechanism of *BRCA1* inactivation that is present in approximately 11% of ovarian cancers and 13-25% of breast cancers (Baldwin et al., 2000; Bell et al., 2011; Esteller et al., 2000; Y. Xu et al., 2013). Lastly, proper gene selection for panel-based tests is limited by our incomplete understanding of non-BRCA causes of HRD. Given the constraints of genetic testing, alternative HRD detection methods are gaining traction. Quantification of RAD51 foci, which mark a key step in the HR pathway, provides a relatively direct test of HRD. However, clinical implementation of RAD51 foci quantification and other functional assays remains challenging, as these assays are labor-intensive and often require fresh tumor tissue (Wijk et al., 2022). Moreover, the RAD51 foci assay is insensitive to downstream defects in HR such as the loss of RAD51AP1 (Modesti et al., 2007). Another approach for detecting HRD assesses the mutational footprints, or genomic “scars”, characteristic of HRD tumors. Since these genomic scars accumulate as a consequence of HRD, they enable HRD detection regardless of the cause. Genomic scar-based assays are also clinically amenable and have been approved as diagnostic tests for PARP inhibitor therapy in ovarian cancer patients (Ngoi & Tan, 2021).

Multiple methods are available for predicting HRD based on genomic scars (Table 1). One of the most commonly used metrics is the “HRD score”, which represents the sum of the counts of three different types of large-scale changes in allelic content/copy number that have been shown to associate with *BRCA1/2* deficiency: Loss of Heterozygosity (LOH), Large-scale State Transitions (LST), and telomeric allelic imbalance (TAI) (Abkevich et al., 2012; Birkbak et al., 2012; Popova et al., 2012; Telli et al., 2016; Timms et al., 2014). To clearly distinguish HRD scores from other HRD metrics, we subsequently refer to HRD scores as “HRDsum” scores. More recently, machine-learning approaches such as HRDetect and Classifier of HOmologous Recombination Deficiency (CHORD) have identified additional combinations of genomic scars predictive of BRCA1/2 deficiency (Davies et al., 2017; L. Nguyen et al., 2020). CHORD primarily uses the relative counts of microhomology-mediated deletions and 1-10 kb duplications to predict HRD, whereas HRDetect combines the proportion of microhomology-mediated deletions with the HRDsum score and four different mutational signature scores to predict HRD. Compared to HRDetect, CHORD requires less pre-processing of the data, was trained on a pan-cancer cohort as opposed to a breast cancer cohort and includes the ability to predict *BRCA1* vs *BRCA2* subtypes (Table 1). HRDetect and CHORD both perform better than HRDsum scores at predicting BRCA1/2 deficiency, but HRDetect and CHORD were designed for analysis of whole-genome sequencing (WGS), whereas HRDsum scores can be generated from more readily available data types, including SNP arrays and whole-exome sequencing (WES) (Table 1).

**Table 1.**
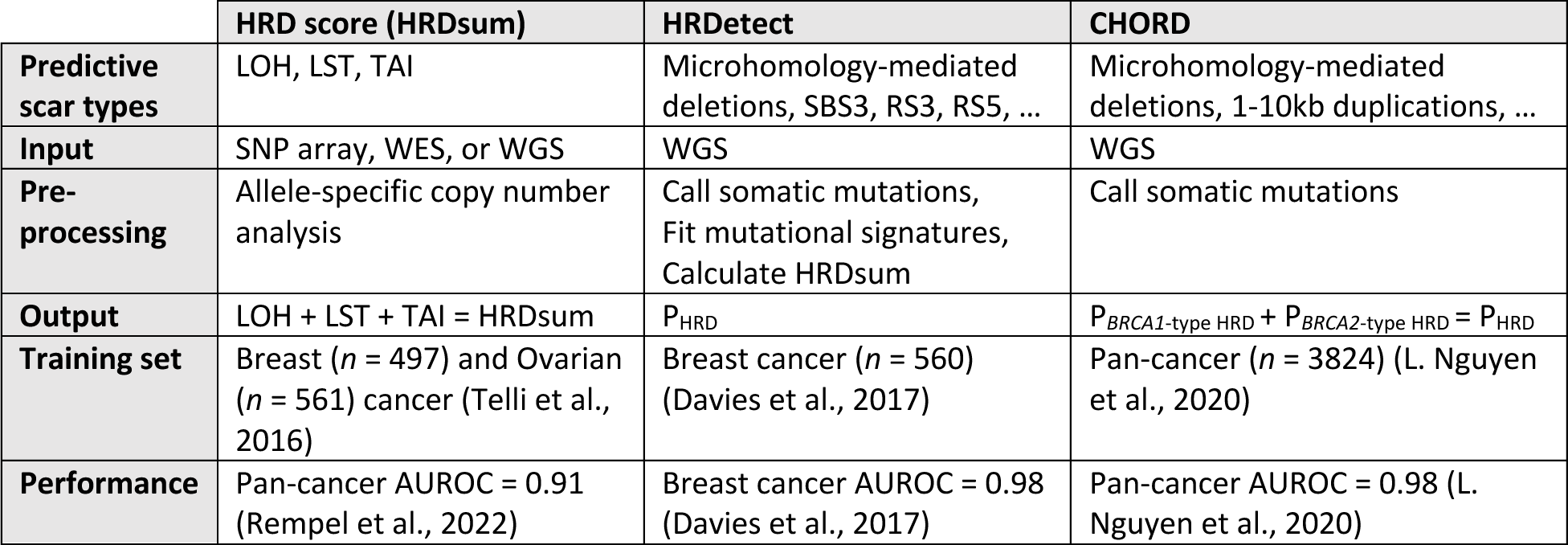
Comparison of common HRD detection methods based on genomic scars. Only the top predictive scar types are shown for HRDetect and CHORD. Calling somatic mutations in preparation for HRDetect and CHORD includes single nucleotide variants, small indels, and structural variants. Performance is represented as the Area Under the Receiver Operating Characteristic curve (AUROC) for predicting BRCA1/2 deficiency. LOH, Loss of Heterozygosity; LST, Large-scale State Transition; TAI, Telomeric Allelic Imbalance; SBS, Single Base Substitution Signature; RS, Rearrangement Signature; SNP, Single Nucleotide Polymorphism, WES, Whole-Exome Sequencing; WGS, Whole-Genome Sequencing; P_HRD_, HRD probability; P*_BRCA1_*_-type HRD_, *BRCA1*-type HRD probability; P*_BRCA2_*_-type HRD_, *BRCA2*-type HRD probability.

|  | HRD score (HRDsum) | HRDetect | CHORD |
| --- | --- | --- | --- |
| **Predictive scar types** | LOH, LST, TAI | Microhomology-mediated deletions, SBS3, RS3, RS5, ... | Microhomology-mediated deletions, 1-10kb duplications, ... |
| **Input** | SNP array, WES, or WGS | WGS | WGS |
| **Pre-processing** | Allele-specific copy number analysis | Call somatic mutations, Fit mutational signatures, Calculate HRDsum | Call somatic mutations |
| **Output** | LOH + LST + TAI = HRDsum | $P_{HRD}$ | $P_{BRCA1\text{-type HRD}} + P_{BRCA2\text{-type HRD}} = P_{HRD}$ |
| **Training set** | Breast ($n = 497$) and Ovarian ($n = 561$) cancer (Telli et al., 2016) | Breast cancer ($n = 560$) (Davies et al., 2017) | Pan-cancer ($n = 3824$) (L. Nguyen et al., 2020) |
| **Performance** | Pan-cancer AUROC = 0.91 (Rempel et al., 2022) | Breast cancer AUROC = 0.98 (Davies et al., 2017) | Pan-cancer AUROC = 0.98 (L. Nguyen et al., 2020) |

Genomic scar-based HRD predictions are available for thousands of patient tumor samples, thus enabling the identification of associations between HRD and other features such as gene deficiencies, cancer types, and clinical outcomes (Davies et al., 2017; Knijnenburg et al., 2018; Marquard et al., 2015; L. Nguyen et al., 2020; Rempel et al., 2022). Given that a wide range of datasets and experimental setups are available exclusively to cell lines, HRD predictions for cell lines could provide additional insights into the molecular underpinnings of its mechanism. To this end, we adapted the CHORD method to cell lines to generate HRD predictions for over 300 cancer cell lines. To predict HRD in cell lines that lacked WGS and were therefore ineligible for CHORD analysis, we also calculated HRDsum scores, resulting in HRD predictions for a total of 1,416 cell lines. We validated the quality of these predictions by comparing them to known *BRCA1/2* statuses and demonstrate the utility of these datasets by using them to explore associations between HRD and various other features in cell lines.

## MATERIALS AND METHODS

### Whole-genome sequencing for CHORD

CHORD was applied to 336 cell lines with WGS available. FASTQ files for 329 of these cell lines were obtained from the Cancer Cell Line Encyclopedia (CCLE) (Ghandi et al., 2019). The other seven cell lines were sequenced for this study as follows: 647-V (DSMZ), J82 (ATCC), JIMT1 (DSMZ), MDA-MB-436 (ATCC), NCI-H1915 (ATCC), SUSA (DSMZ), and UWB1.289 (ATCC) cells were cultured for 48-72 hours at 37°C, 5% CO_2_ in the supplier’s recommended media and collected for gDNA isolation. gDNA was isolated on an XTRACT 16+ (AutoGen) using an XK110-96 kit and quantified on a Qubit 4 (Invitrogen). Samples were submitted to the Broad Clinical Research Sequencing Platform for PCR-free 30x WGS. For all samples, reads were aligned to human genome reference GRCh38 using BWA-MEM (v0.7.17) (H. Li & Durbin, 2009).

### Variant calling and filtering for CHORD

Prior to calling variants, duplicate reads were marked and removed using the MarkDuplicates algorithm from GATK (v4.1.7.0). Single nucleotide variants (SNV) and small insertions and deletions (indels) were detected using SAGE (v3.0.1) and annotated using PAVE (v1.2). SAGE and PAVE are both available from the Hartwig Medical Foundation (HMF, https://github.com/hartwigmedical/hmftools). Since SAGE is optimized for samples with 100x coverage by default, we made the following adjustments for analysis of the 30x WGS samples: min_tumor_qual filters were adjusted to the cutoffs recommended for 30x coverage (Hotspot = 40; Panel = 60; High Confidence = 100; Low Confidence = 150), and the read depth parameters max_read_depth, max_read_depth_panel, and max_realignment_depth were lowered to 400, 40,000, and 400, respectively. Using the GRCh38 HMF cohort panel of normals as a reference, common variants were removed with the recommended cut-offs (HOTSPOT:5:5;PANEL:2:5;UNKNOWN:2:0) during the PAVE annotation step. Structural variants were detected using GRIDSS (v2.13.2) (Cameron et al., 2017, 2021). The HMF tool GRIPSS (https://github.com/hartwigmedical/hmftools) was used to remove low-quality structural variants and flag structural variants present in HMF cohort panels of normals.

In addition to removing variants marked as “PON” (Panel Of Normals) by PAVE and GRIPSS, we applied a variety of other filters to help select for high-quality somatic variants. First, indels were normalized, multi-nucleotide variants were split into SNVs, and multiallelic SNVs and indels were separated into multiple entries. Then, SNVs and indels present in gnomAD v2.1, dbSNP build 146, and/or the Homo_sapiens_assembly38.known_indels.vcf file from the GATK resource bundle were removed. SNVs, indels, and structural variants were also removed if they were present in 3 or more cell lines, a cut-off that was chosen based on CHORD performance. To obtain high-quality variants, we selected PASS variants with allele frequencies ≥ 0.1. For SNVs and indels, we also required that variants in low-confidence regions have a quality score ≥ 240 and variants in high-confidence regions have a quality score ≥ 170. For structural variants, we required that variants have a quality score ≥ 1000 and have supporting assemblies from both sides of the breakpoint (AS > 0 & RAS > 0). All variant processing and filtering steps were performed using BCFtools (Danecek et al., 2021).

### CHORD analysis

In preparation for CHORD, structural variant lengths and categories (deletions, duplications, inversions, and translocations) were determined using code adapted from https://github.com/PapenfussLab/gridss/blob/master/example/simple-event-annotation.R. Mutation “contexts”, or counts for various mutation types, were extracted and then used to generate CHORD predictions with the CHORD package in R (Supplementary Table S1) (L. N. Nguyen, 2022). In the original CHORD analysis of patient samples, tumors containing more than 14,000 indels in repeat regions were classified as showing MicroSatellite Instability (MSI) (L. Nguyen et al., 2020). For cell lines, we found that a cut-off of 10,000 indels in repeat regions resulted in better alignment with previously determined MSI annotations (Chan et al., 2019). Therefore, the min.msi.indel.rep parameter was lowered to 10,000. All other parameters remained unchanged from the default setting.

### Gene deficiency and proficiency labeling

*BRCA1*, *BRCA2*, and other genes included in this study were labeled as deficient if they met one of the two following criteria: 1) The gene was homozygous (allele frequency > 0.85) for a mutation that was annotated as pathogenic by ClinVar and/or predicted to have a high impact by VEP (v106) (McLaren et al., 2016). For this analysis, cell line mutations were obtained from the DepMap portal (https://depmap.org) under release 22Q4 and then annotated with VEP. 2) The gene contained a deep deletion, as determined by copy number analysis (WES_pureCN_CNV_genes_cn_category_20221213.csv) from the Sanger Institute (https://cellmodelpassports.sanger.ac.uk/downloads). For *BRCA1* and *BRCA2*, we also reviewed the literature for additional information on functional status. With this approach, we identified HCC1428 as a *BRCA2* revertant.

For assessment of CHORD performance, BRCA1/2 deficiency and proficiency labels were assigned as follows. Cell lines that met one of the deficiency criteria described above were labeled as BRCA1/2-deficient. Cell lines were labeled as BRCA1/2-proficient if they: 1) lacked any mutations in *BRCA1* and *BRCA2*, according to the mutation calls from DepMap (22Q4); 2) lacked any deep deletions in *BRCA1* and *BRCA2*, according to the copy number calls from the Sanger Institute; and 3) expressed *BRCA1* and *BRCA2* at levels above the lower quartile (log2(TPM+1) = 3.8 for *BRCA1*, 2.07 for *BRCA2*). Expression levels were obtained from the DepMap portal (https://depmap.org) under release 22Q4.

SIFT and PolyPhen predictions for the *BRCA1* variant Y1853C were determined using VEP (v106) (McLaren et al., 2016).

To identify cases of *BRCA1* gene silencing, we looked for cell lines that showed both relatively low expression of *BRCA1* and relatively high levels of *BRCA1/NBR2* promoter methylation. Expression values were downloaded from the DepMap portal (release 22Q4), and promoter methylation values were obtained from the 2019 CCLE (Ghandi et al., 2019). We defined low *BRCA1* expression as log2(TPM+1) values below the lower quartile (3.8) and high promoter methylation as methylation fraction values > 0.3. When comparing *BRCA1* expression to *NBR2* promoter methylation, a defined cluster of cell lines (MDAMB134VI, EFO21, TE5, OVCAR4, SNU119, OVCAR8, and CAL851) met these criteria. OVCAR8 was also reported in the literature as showing *BRCA1* methylation (Stordal et al., 2013). Promoter methylation data for HCC38 was unavailable from the CCLE. However, HCC38 was previously reported as showing *BRCA1* promoter hypermethylation and minimal *BRCA1* expression (Stefansson et al., 2012). Therefore, we annotated HCC38 as a likely case of *BRCA1* gene silencing, along with the aforementioned cell lines.

### HRDsum score analysis

HRDsum scores were determined from three different allele-specific copy number datasets. The first dataset, referred to as the “Broad” dataset, was acquired from the 2019 CCLE study and contains copy number calls for 997 cell lines analyzed by the Broad Institute (Ghandi et al., 2019). The second and third datasets, referred to as “Sanger (Broad WES)” and “Sanger (Sanger WES)”, were both derived from copy number calls (WES_pureCN_CNV_segments_20220623.csv) provided by the Sanger Institute (https://cellmodelpassports.sanger.ac.uk/downloads). The “Sanger (Broad WES)” dataset contains copy number calls for 324 cell lines with WES originating from the Broad Institute, and the “Sanger (Sanger WES)” dataset contains copy number calls for 1,047 cell lines with WES originating from the Sanger Institute. HRDsum scores were calculated from each dataset using code adapted from a previous study (Marquard et al., 2015).

After generating HRDsum scores for each dataset, we merged the three sets of scores into one summary dataset. First, we calculated the median HRDsum score for each of the 104 cell lines that were shared by all three datasets. Then, for each dataset, we fit a regression model using natural splines to predict the median HRDsum score for the 104 common cell lines. The three resulting models were applied to their corresponding datasets to generate normalized HRDsum scores for all cell lines. The normalized scores from all three datasets were combined, and the mean score was calculated for cell lines present in more than one dataset. These “summary” HRDsum scores were used throughout this study and are available as a supplemental table, along with the raw HRDsum scores for each dataset.

To assign tissue types to each cell line in the HRDsum score dataset, we harmonized the Broad Institute and Sanger Institute tissue type annotations. Broad Institute annotations were downloaded from the DepMap portal (https://depmap.org) under release 22Q4, and Sanger Institute annotations (model_list_20230110.csv) were downloaded from https://cellmodelpassports.sanger.ac.uk/downloads. Sanger tissue terms were converted to Broad tissue terms, and Broad annotations were used in cases where the two different annotations did not agree. Non-cancerous cell lines were grouped into a separate category labeled “Non-Cancerous”. These tissue annotations were used throughout this study.

### Mutation enrichment analysis

To test if HRD cell lines were enriched for deficiencies in any DNA repair-related genes, we first obtained a list of 276 genes involved in DNA damage repair that was curated in a previous study (Knijnenburg et al., 2018). Deficiencies in these genes were determined as described above. Enrichment was determined with a one-tailed Fisher’s Exact test, and *p*-values were adjusted with the Benjamini-Hochberg method. Only genes showing a deficiency in at least one cell line classified as HRD were included in the analysis.

### Association analyses

Associations between HRD predictions and other features were determined with either a Pearson’s correlation test or by binning cell lines based on HRDsum quartiles and performing a Kruskal-Wallis test. For analyses exploring associations among multiple CCLE features or drug sensitivities, *p*-values were adjusted with the Benjamini-Hochberg method. Non-cancerous and MSI cell lines were excluded from these analyses. Gene expression values, damaging mutations, relative copy number values, and gene dependency scores were downloaded from the DepMap portal (https://depmap.org) under release 22Q4. Promoter methylation fractions were obtained from the 2019 CCLE study (Ghandi et al., 2019). For the PRISM Repurposing drug response dataset, which was downloaded from the DepMap portal under release 23Q2, associations were tested using the log2 fold change in viability (treatment versus DMSO) (Corsello et al., 2020). For the GDSC2 drug response dataset, the July 24, 2022 release was downloaded from https://www.cancerrxgene.org/downloads, and associations were tested using the log10 IC_50_ values (Yang et al., 2013). Annotations for MSI and p53 functional status were obtained from a previous study (Chan et al., 2019).

### PARPi Sensitivity Determination

Sensitivity to Niraparib or Olaparib was measured via clonogenic assays. Cells were seeded in 6-well dishes in the supplier’s recommended media and incubated at 37°C, 5% CO_2_ overnight. The following day, Niraparib (SelleckChem) or Olaparib (SelleckChem) was added at a top dose of 10µM across a 10-point, 1:3 dilution curve. Following 7 days of drug treatment, media was removed and fresh media containing drug was added. On day 14, media was removed and cells were fixed with 4% paraformaldehyde for 15 minutes. Cells were washed with water and stained with 0.1% crystal violet in a 10% (v/v) ethanol in dH_2_O solution for 20 minutes. Plates were washed with water and dried overnight at room temperature. Crystal violet was extracted using 10% acetic acid and absorbance was read using a BioTek Cytation 5 plate reader.

Absorbance values were used to quantify IC_50_ values via the following procedure: For each assay, a spreadsheet containing absorbance values was ingested into an internal LIMS database using the Apache POI library (version 4.1.1) and values were normalized to the average value of the wells containing the control (e.g. DMSO). From these normalized values, single agent IC_50_ values were calculated by fitting the single agent data to a two-parameter dose response curve of the form y = 1 −1 / (1 + 10^(10^b * (x -a))), where y was the response, with 0 indicating no response and 1 indicating complete inhibition; x was the log10 of the concentration in nM; and the IC_50_ value was given by 10^a. If the optimization function failed to find a fit using this model, fitting was performed using the alternative model y = 1 / (1 + e^(-|b| * (x -a))), where x and y served the same roles as previously described, and where the IC_50_ value was given instead by a. In both cases, fitting was performed by the SimpleCurveFitter.fit method of the Apache Commons Math library (version 3.6.1).

Cell lines were classified as “sensitive” to PARP inhibition if at least 66% of replicates showed an IC_50_ value < 1 μM and a maximal inhibitory response > 75%. Cell lines were classified as “insensitive” to PARP inhibition if at least 66% of replicates showed an IC50 value ≥ 1 μM and a maximal inhibitory response < 35%. Cell lines that did not meet either the “sensitive” or “insensitive” criteria were excluded from the analysis.

### Statistical analysis

All statistical analyses were performed using R version 4.1.3 (R_Core_Team, 2022). Specific statistical tests along with *p*- or *q*-values are described in each figure legend.

### Data availability

WGS data generated for this study are archived in GEO under accession number XXXX. All other data generated in this study are available in the paper and supplemental tables. Sources for publicly available datasets analyzed in this study are described above. Source code for the generation of CHORD probabilities and HRDsum scores is available at https://github.com/ksqtx/XXXX.

## RESULTS

### CHORD performance in cell lines

To predict HRD in cell lines, we first applied the CHORD model to 336 cell lines with WGS available (Fig. 1a). CHORD predicts HRD based on somatic variants, which can be called by comparing variants between tumor and matching normal samples. However, matching normal samples are not available for most cancer cell lines. To identify probable somatic variants for cell lines without matching normal samples, we inferred variants in tumor-only mode and then applied multiple filters that remove known germline variants as well as variants that are likely artifacts of PCR amplification or sequencing (Fig. 1a, see “Methods”). Since CHORD relies on relative counts of microhomology-mediated deletions and other mutation types, it cannot be applied to samples with MSI, as these samples contain an abundance of indels in repeat regions that dominate the signal (L. Nguyen et al., 2020). CHORD classified 32 cell lines as MSI, all of which were called as MSI in an independent study (Supplementary Fig. S1) (Chan et al., 2019). For the remaining 304 cell lines, CHORD generated probability scores for both *BRCA1*-type and *BRCA2*-type HRD as well as an overall HRD probability, which is equal to the sum of the two subtype scores (Supplementary Table S2).

**Figure 1.**
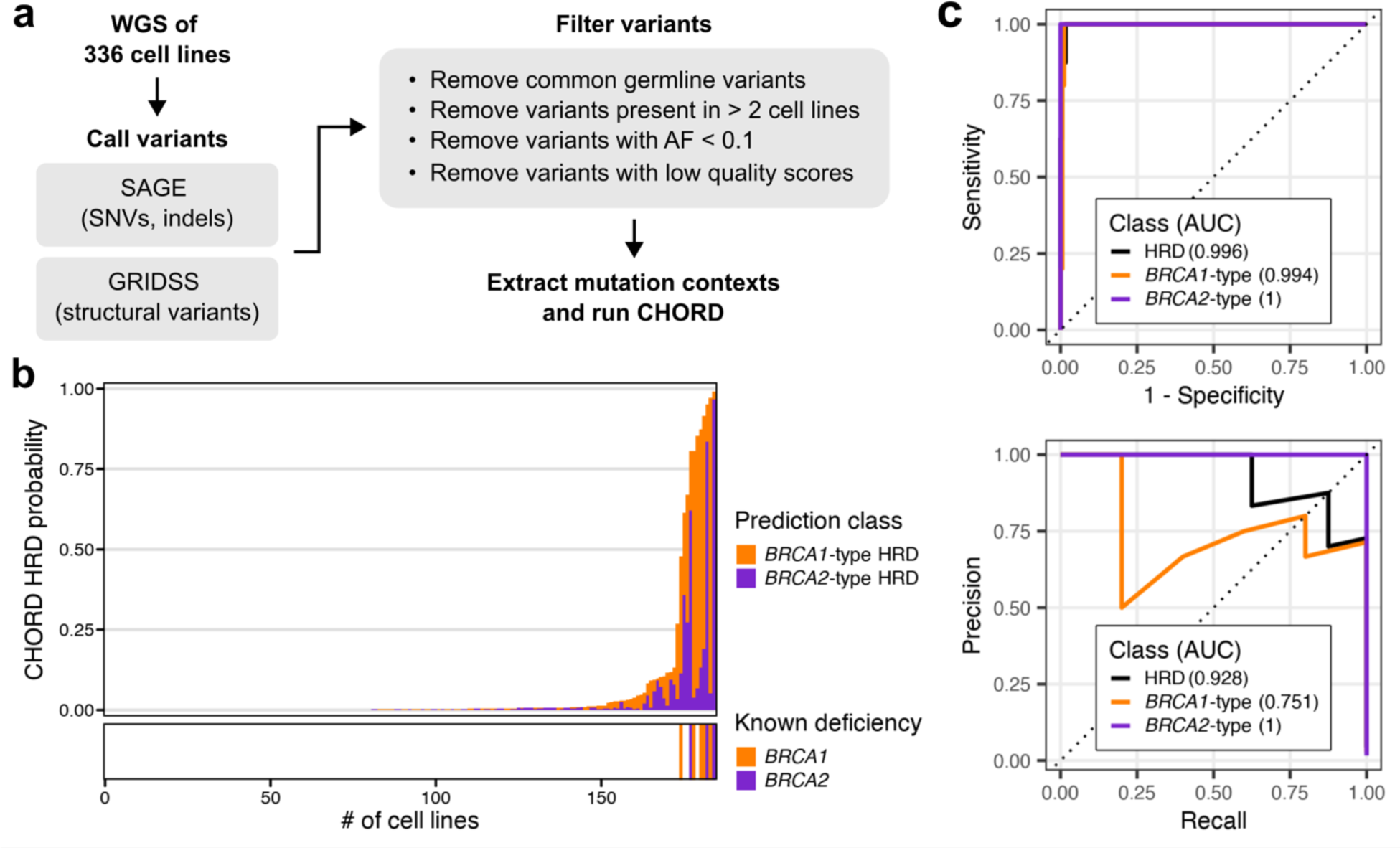
Adapting CHORD for cell lines. **a)** Pre-processing steps to prepare data for CHORD. For cell lines with WGS available, single nucleotide variants and small indels were called using SAGE, and structural variants were called using GRIDSS. Filters were applied to select for quality variants that are likely somatic prior to running CHORD. AF, Allele Frequency. **b)** CHORD predictions for BRCA1/2-deficient and likely BRCA1/2-proficient cell lines. Top panel: CHORD HRD probabilities shown as the sum of *BRCA1*-type and *BRCA2*-type probabilities. Bottom panel: Cell lines with biallelic loss of *BRCA1* or *BRCA2*. **c)** CHORD performance as determined by the Receiver Operating Characteristic (top) and Precision-Recall (bottom) curves for HRD, *BRCA1*-type, and *BRCA2*-type prediction classes. Area under the curve (AUC) is shown in parentheses.

To test how well CHORD performs on cell lines, we compared the subtype and overall HRD probabilities of cell lines showing biallelic loss of *BRCA1* or *BRCA2* (*n* = 8) to the corresponding probabilities of cell lines likely to be proficient for BRCA1/2 (*n* = 176). BRCA1/2 proficiency was defined as cell lines that are wild-type for *BRCA1* and *BRCA2* and also show *BRCA1/2* expression levels above the lower quartile. Overall, CHORD predictions agreed well with the known BRCA status of these cell lines, in terms of both the overall HRD probabilities as well as the *BRCA1/2* subtype predictions (Fig. 1b). To quantify CHORD performance, we first measured the Area Under the Receiver Operating Characteristic curve (AUROC). The AUROC values were 0.996, 0.994 and 1 for overall HRD predictions, *BRCA1*-type predictions and *BRCA2*-type predictions, respectively (Fig. 1c). Given the class imbalance (8 BRCA1/2-deficient versus 176 BRCA1/2-proficient cell lines), we also measured the Area Under the Precision-Recall curve (AUPRC). The AUPRC values were 0.928, 0.751 and 1 for overall HRD predictions, *BRCA1*-type predictions and *BRCA2*-type predictions, respectively (Fig. 1c). Since all known BRCA1/2-deficient cell lines showed an HRD probability > 0.4, we set this value as the cut-off for classification and subsequently refer to cell lines with HRD probabilities > 0.4 as CHORD-HRD and cell lines with HRD probabilities ≤ 0.4 as CHORD-HR-proficient, or CHORD-HRP. With an HRD probability cut-off of 0.4, three of the 11 cell lines predicted to be HRD (CAOV3, PANC0327 and UACC893) were initially labeled as BRCA1/2-proficient, as they lack detectable mutations in *BRCA1/2* and show intermediate to high levels of *BRCA1/2* expression (Supplementary Table S3). It is possible that these three cell lines are indeed false positives. However, given that CHORD is intended to capture instances of HRD that are overlooked by more traditional detection methods, it is also possible that these cell lines are truly HRD, perhaps due to non-BRCA causes that would have escaped detection by our labeling criteria. Potential causes of HRD in these cell lines are further explored in the following section.

### Genetic and epigenetic *BRCA1/2* alterations explain the majority of CHORD HRD classifications

After assessing the performance of CHORD on BRCA1/2-deficient and likely BRCA1/2-proficient cell lines, we extended our analysis to all 336 cell lines with WGS available. In addition to the 8 cell lines known to be BRCA1/2-deficient, 12 other cell lines were also classified as CHORD-HRD (Fig. 2a). This included KE39, a gastric carcinoma cell line that is homozygous for a missense mutation in *BRCA1* (Y1853C) but was not originally labeled as BRCA1/2-deficient since this mutation was not deemed damaging by our initial criteria (see “Methods”). To further assess the Y1853C variant, we predicted its functional impact using SIFT and PolyPhen and found that Y1853C scored as “deleterious” with a value of 0 and as “probably damaging” with a value of 1, respectively. Furthermore, the consensus for Y1853C in the ClinVar database is “likely pathogenic”. These observations, combined with the *BRCA1*-type HRD prediction made by CHORD, suggest that KE39 is deficient for BRCA1. Since genomic scars serve as a historical record of repair deficiencies and do not necessarily reflect the current repair status, CHORD and other scar-based methods are likely to classify *BRCA1/2*-revertant lines as HRD. Indeed, HCC1428, a *BRCA2*-mutated cell line with a secondary mutation that rescues BRCA2 function, showed an HRD probability of 0.792 (Fig. 2a).

**Figure 2.**
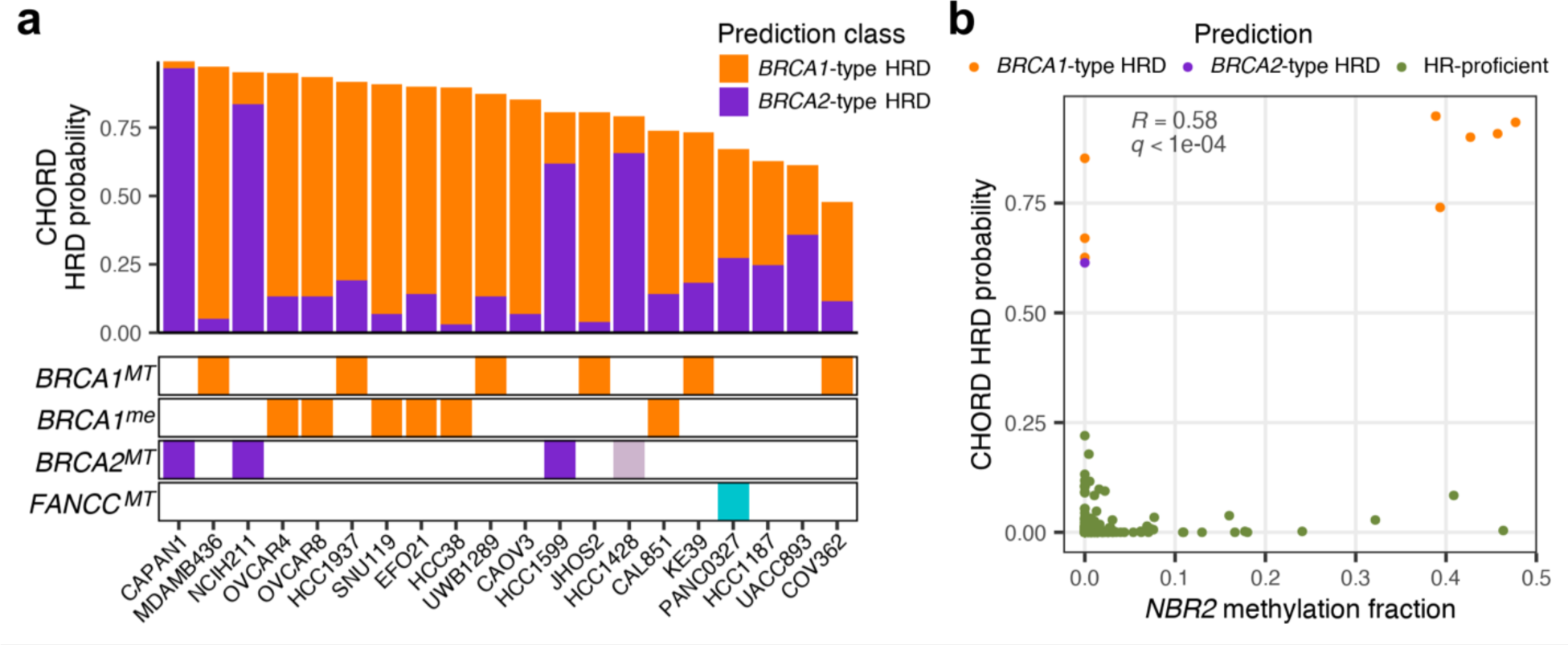
HR gene alterations in CHORD-HRD cell lines. **a)** Potential causes of HRD in CHORD-HRD cell lines. Top panel: CHORD HRD probabilities shown as the sum of *BRCA1*-type and *BRCA2*-type probabilities. Bottom panels: Colored bars indicate biallelic loss of *BRCA1*, *BRCA2*, or *FANCC* (*BRCA1^MT^*, *BRCA2^MT^*, *FANCC^MT^*) or promoter hypermethylation of *BRCA1* (*BRCA1^me^*). The lighter *BRCA2^MT^* color for HCC1428 represents the presence of a secondary mutation. **b)** Correlation between CHORD HRD probabilities and *NBR2* promoter methylation levels in wild-type *BRCA1/2* cell lines. Colors represent CHORD classifications. Pearson correlation coefficient (*R*) and Benjamini-Hochberg-adjusted *p*-value (*q*) are shown in gray.

Genetic inactivation of *BRCA1/2* was not detected in the remaining 10 cell lines classified as HRD. To explore the underlying causes of HRD in these cell lines, we calculated Pearson correlations between CHORD HRD probabilities and various features available from the CCLE in wild-type *BRCA1/2* cell lines. The most significant association was a positive correlation with promoter methylation of *NBR2* (*neighbor of BRCA1 gene 2*), with five *BRCA1*-type CHORD-HRD lines showing relatively high levels of *NBR2* promoter methylation (*q* < 0.0001; Fig. 2b). Interestingly, *NBR2* shares a bidirectional promoter with *BRCA1* (C.-F. Xu et al., 1997). CHORD HRD probabilities also correlated positively with *BRCA1* promoter methylation levels (*q* < 0.0001 for sites chr17:41276132-41277132 and chr17:41277500-41278500) and negatively with *BRCA1* expression levels (*q* < 0.0001) (Supplementary Fig. S2a). Together, these observations suggest that multiple *BRCA1*-type CHORD-HRD lines are HRD due to epigenetic silencing of *BRCA1*. To identify all likely cases of *BRCA1* silencing, we determined which cell lines showed evidence of *BRCA1* silencing in previous studies and/or show both relatively high levels of *NBR2/BRCA1* promoter methylation and relatively low levels of *BRCA1* expression (see “Methods”; Supplementary Fig. S2b). Out of the eight cell lines that met these criteria, six cell lines (CAL851, EFO21, HCC38, OVCAR4, OVCAR8, and SNU119) were classified by CHORD as *BRCA1*-type HRD (Fig. 2a). Thus, epigenetic inactivation of *BRCA1* likely explains a substantial fraction of HRD classifications (6/10) in wild-type *BRCA1/2* cell lines.

After accounting for both the genetic and epigenetic status of *BRCA1/2*, four CHORD-HRD cell lines (CAOV3, PANC0327, HCC1187, and UACC893) lacked any detectable BRCA1/2 alterations. To investigate the potential causes of HRD in these cell lines, we determined if biallelic loss of any DNA repair-related genes other than *BRCA1/2* was enriched in these cell lines using a previously curated list of genes involved in major DNA damage repair pathways (Knijnenburg et al., 2018). Although no genetic deficiencies were significantly enriched after adjusting for multiple testing, it is interesting to note that the top hit was *FANCC*, which encodes a Fanconi anemia (FA) protein that has been shown to promote homologous recombination (Supplementary Table S4) (Niedzwiedz et al., 2004). The *BRCA1*-type CHORD-HRD cell line PANC0327 harbors a deletion of *FANCC*, whereas none of the CHORD-HRP cell lines showed biallelic inactivation of *FANCC* (Fig. 2a). In patient samples, deficiencies in *RAD51C* and *PALB2* associate with *BRCA2*-type HRD predictions (L. Nguyen et al., 2020). In cell lines, however, associations with these genetic deficiencies could not be tested, as none of the cell lines analyzed by CHORD show biallelic inactivation of either *RAD51C* or *PALB2*. Aside from the *FANCC* deletion in PANC0327, no other DNA repair-related gene deficiencies were exclusive to CHORD-HRD cell lines, and the underlying cause of the HRD phenotype for CAOV3, HCC1187, and UACC893 remains to be determined.

### Generation of HRDsum scores for cell lines

The above results suggest that, with the proper pre-processing of the data, CHORD can be a powerful tool for predicting HRD and *BRCA1/2* subtypes in cell lines. In its current state, however, the CHORD method is only applicable to cell lines with WGS available. In comparison, HRDsum scores can be determined from more readily available types of data such as SNP arrays or WES. Therefore, to gain a more comprehensive understanding of HRD in cell lines, we also calculated HRDsum scores for cell lines using an approach that was previously applied to patient samples (Marquard et al., 2015). Starting with allele-specific copy number calls available from either the Broad or Sanger Institutes, we generated HRDsum scores for three different datasets (Fig. 3a). A subset of cell lines were present in more than one dataset, enabling us to compare HRDsum scores between the different datasets for these cell lines (Fig. 3b). Although there was a shift in absolute values between the datasets, HRDsum scores derived from different data sources were highly correlated, with Pearson correlation coefficient values ≥ 0.89 (Fig. 3c). We used natural spline regression to integrate the different datasets into a single “summary” HRDsum score for each cell line (see “Methods”), resulting in a dataset of HRDsum scores for 1,414 cell lines (Supplementary Table S5).

**Figure 3.**
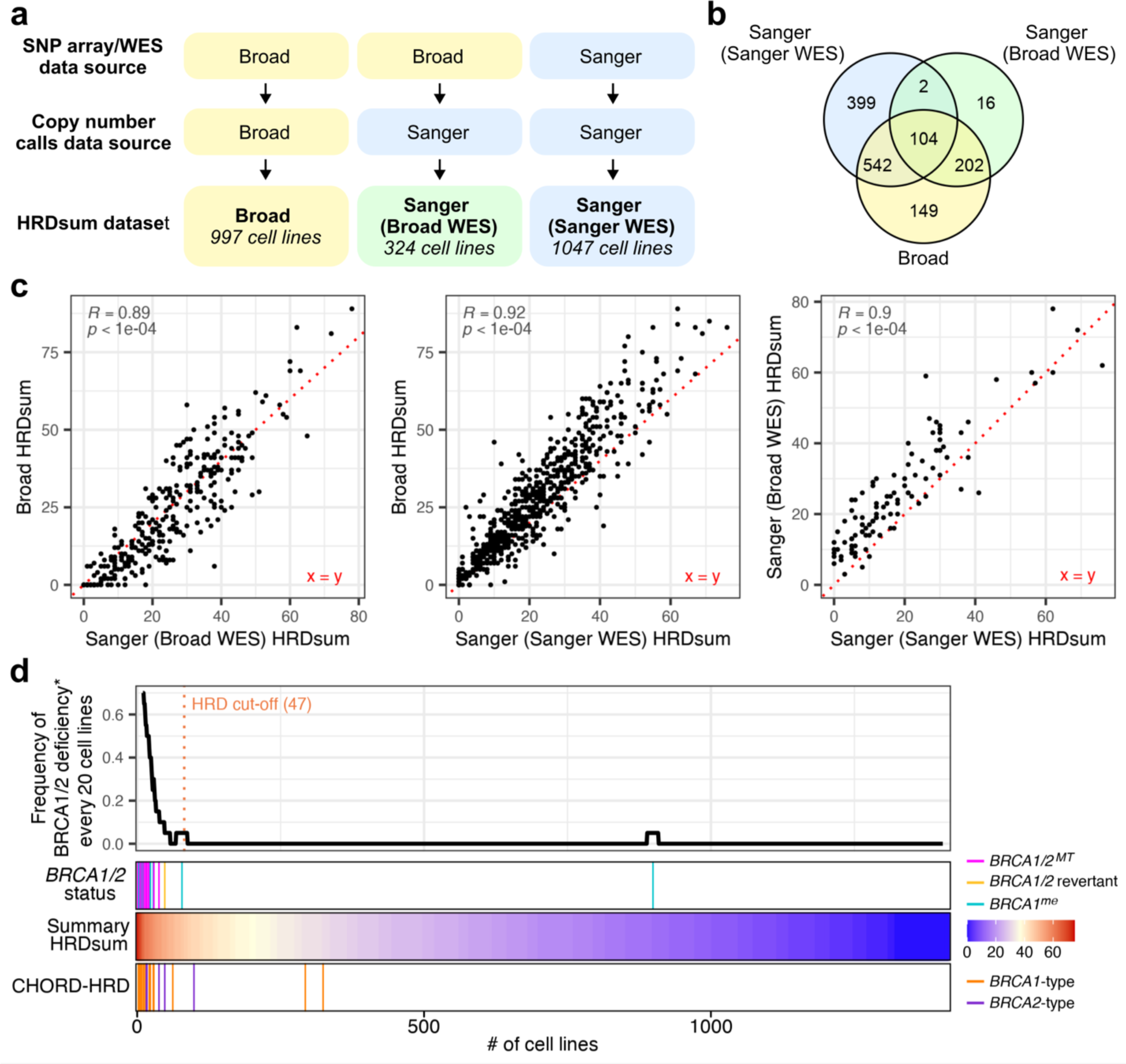
HRDsum scores in cell lines. **a)** HRDsum scores were calculated for three different datasets with different sources of SNP array/WES data and/or copy number calls. **b)** Venn diagram showing the overlap in cell lines between the three different datasets. **c)** Scatter plots comparing HRDsum scores between pairs of datasets for cell lines that were in common. Pearson correlation coefficients (*R*) and *p*-values are shown in gray. **d)** Occurrence of *BRCA1/2* loss and CHORD-HRD predictions in relation to HRDsum scores. Top panel: Rolling frequency of BRCA1/2 deficiencies for every 20 cell lines, with cell lines ranked by HRDsum score. *The term “BRCA1/2 deficiency” includes both current deficiencies (biallelic loss of *BRCA1/2* or likely epigenetic silencing of *BRCA1*) as well as historical deficiencies (*BRCA1/2* reversions). A cut-off (orange line) was set at the 5^th^ percentile of HRDsum scores for cell lines with BRCA1/2 deficiencies. Bottom panels: Colored bars indicate BRCA1/2 deficiency types, summary HRDsum scores, and *BRCA1/2* subtype predictions for CHORD-HRD cell lines, in order from top to bottom.

Like CHORD probabilities, HRDsum scores agreed well with known *BRCA1/2* status. As expected, BRCA1/2-deficient cell lines ranked among the highest HRDsum scores (Fig. 3d). HRDsum scores in cell lines with either biallelic loss of *BRCA1/2* or likely epigenetic silencing of *BRCA1* were significantly higher than HRDsum scores in tissue-matched cell lines that are likely BRCA1/2*-*proficient (Supplementary Fig. S3a). As was the case with CHORD (and is likely the case for most scar-based HRD detection methods), the HRDsum score method produced a relatively high score (52) for the *BRCA2*-revertant cell line HCC1428 (Figure 3d). Overall, the strong association between HRDsum scores and *BRCA1/2* alterations suggests that HRDsum scores serve as an independent and high-quality proxy of HRD status.

For patient samples, analyses of HRDsum scores often apply a cut-off of ≥ 42, which was set by calculating the 5^th^ percentile of HRDsum scores for a group of BRCA1/2-deficient breast and ovarian tumors (Telli et al., 2016). For cell lines, we propose a cut-off of ≥ 47, which was the 5^th^ percentile of HRDsum scores for BRCA1/2-deficient cell lines (Figure 3d). Eighty-two out of 1,414 cell lines (6%) fell above this cut-off and were therefore predicted to be HRD. We subsequently refer to these cell lines as HRDsum-high. BRCA1/2 deficiencies were detectable in 21 (26%) of HRDsum-high cell lines. To explore other possible causes of HRD in the remaining 61 HRDsum-high cell lines, we tested if these cell lines were enriched for biallelic mutations in any other DNA repair-related genes, similar to the analysis performed for the CHORD dataset. The top hit was a deficiency for *ATRX*, which encodes a chromatin remodeler known to function in HR (Supplementary Fig. S3b, Supplementary Table S6) (Juhász et al., 2018). However, the enrichment for *ATRX* mutations was not significant after correcting for multiple testing. In our enrichment analysis for CHORD, we identified *FANCC* loss as a possible explanation for the high HRD probability observed in PANC0327 (Fig. 2a). PANC0327 was not HRDsum-high; however, a second *FANCC*-deficient cell line, HUH7, showed a high HRDsum score of 52 (Supplementary Fig. S3b). HUH7 was not included in the CHORD dataset and therefore the CHORD probability for this cell line remains to be determined.

Next, we compared HRDsum scores to CHORD probabilities for the 302 cell lines that were in common between the two datasets (Supplementary Fig. S3c). CHORD-HRD cell lines showed significantly higher HRDsum scores than CHORD-HRP cell lines, and most (17/20) CHORD-HRD cell lines were HRDsum-high (Fig 3d, Supplementary Fig. S3d). Although the strongest HRD predictions were shared by both the CHORD and HRDsum score datasets, some HRD predictions were unique to one of the two datasets (Supplementary Fig. S3e). UACC893, HCC1187, and PANC0327, for instance, were all classified as CHORD-HRD, but the HRDsum scores for these cell lines fell below the HRDsum cut-off (Fig. 2a, Supplementary Fig. S3e).

### HRD cell lines are primarily breast and ovarian cancer types

Using both the CHORD and HRDsum score datasets, we investigated the prevalence of HRD across different cancer types. In patient samples, CHORD HRD classifications are most frequent in ovarian (30-52%) and breast (12-24%) cancer types, followed by pancreatic (7-13%) and prostate (6-13%) cancers (L. Nguyen et al., 2020). We grouped cell lines by tissue type and observed that, similar to patient samples, CHORD HRD predictions were most frequent in cell lines derived from ovarian (38%) and breast (23%) cancers (Fig. 4a). HRD predictions were also present in cell lines derived from pancreatic (18%), esophagus/stomach (4%), and lung (1%) cancers (Fig. 4a). Similar to the CHORD dataset, the HRDsum score dataset showed the highest prevalence of HRD in breast (33%) and ovarian (21%) tissue types (Fig. 4b). The highest HRDsum score was in the cervical cancer cell line DOTC24510, which is homozygous for a stop-gain variant in *BRCA2*. As expected, non-cancerous cell lines showed the lowest mean HRDsum score (Fig. 4b). The relatively high mean HRDsum score for the Esophagus/Stomach tissue type was somewhat surprising, although a previous pan-cancer analysis of HRDsum scores in patient samples also showed an enrichment of HRD in esophageal carcinomas (Fig. 4b) (Rempel et al., 2022). Although up to 13% of prostate tumor patient samples score as HRD, none of the prostate cancer cell lines included in our analysis qualified as HRD (L. Nguyen et al., 2020; Rempel et al., 2022). However, the CHORD and HRDsum datasets included only 1 and 8 prostate cell lines respectively, and four of those cell lines are MSI, which has been shown to be mutually exclusive with HRD (Budczies et al., 2022; Marquard et al., 2015). Additional samples are needed to obtain a better understanding of HRD in prostate cancer cell lines.

**Figure 4.**
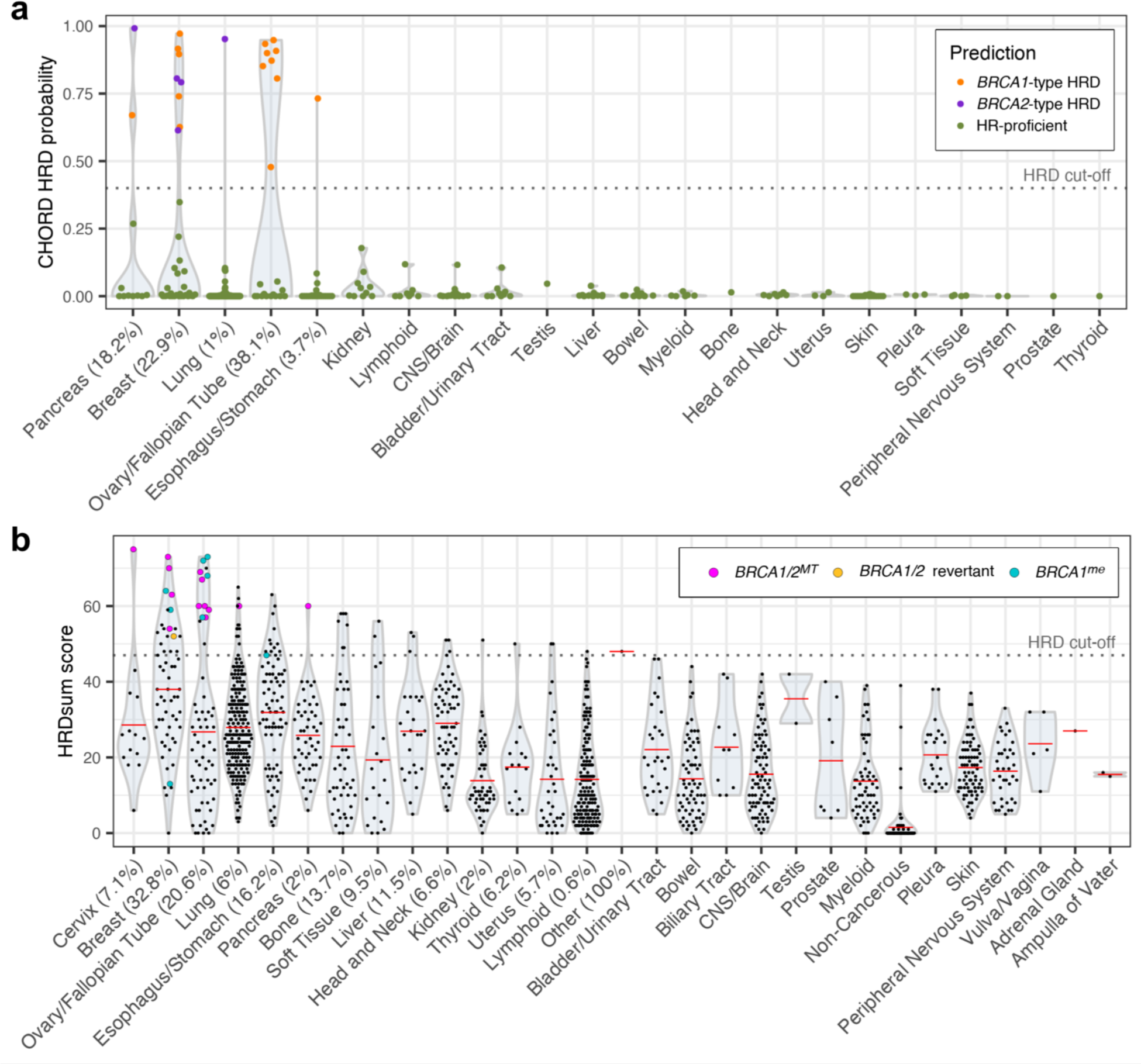
HRD predictions by tissue type. **a)** CHORD HRD probabilities (y-axis) and classifications (color) for cell lines grouped by tissue type. Percentages indicate the percent of cell lines in each tissue type that were classified as HRD using a cut-off of > 0.4 (dashed line). **b)** Violin plot of HRDsum scores in cell lines grouped by tissue type. Non-cancerous cell lines were grouped into a separate “Non-Cancerous” category. Cell lines with biallelic loss of *BRCA1/2*, *BRCA1/2* reversions, or likely epigenetic silencing of *BRCA1* are highlighted as colored points. Percentages indicate the percent of cell lines in each tissue type that were classified as HRD using a cut-off of ≥ 47 (dashed line). Red bars represent the mean HRDsum score.

To further assess HRD by indication, we aligned cell lines with HRDsum scores to transcriptionally similar tumor samples using Celligner (Warren et al., 2021). HRDsum-high cell lines predominantly clustered with tumor samples of the same cancer type, with several cell lines aligning with breast and ovarian tumors, followed by lung and esophageal tumors (Supplementary Fig. S4a). Since CHORD provides subtype-level resolution for HRD, we also mapped CHORD predictions to tumor types to see how the distribution of *BRCA1*-type HRD compared to that of *BRCA2*-type HRD. Interestingly, *BRCA1*-type breast cancer lines all clustered with the basal subtype, whereas *BRCA2*-type breast cancer lines represented a variety of subtypes (Supplementary Fig. S4b). This observation aligns with previous studies showing that patient tumors with germline mutations in *BRCA1*, but not *BRCA2*, express a basal-like phenotype (Foulkes et al., 2003; Lakhani et al., 2005). Overall, these results suggest that the distribution of HRD across different cancer types and subtypes in cell lines is representative of that in patient samples.

### HRDsum scores associated with p53 loss, microsatellite stability, and chromosome 3q26 copy number

In patient samples, HRD exhibits a positive correlation with *TP53* inactivation and an inverse correlation with MSI (Budczies et al., 2022; Knijnenburg et al., 2018; Marquard et al., 2015; Rempel et al., 2022; Takamatsu et al., 2022). To test whether these associations are conserved in cell lines, we first compared HRDsum scores to the functional status of p53 using previously determined predictions based on nutlin-3 sensitivity as well as a p53 target gene expression signature (Chan et al., 2019). Pan-cancer, HRDsum scores were significantly higher in p53-deficient cell lines than in p53-proficient cell lines (Fig. 5a). At the level of individual tissue types, HRDsum scores associated significantly with *TP53* status in breast and esophagus/stomach cancer cell lines (Supplementary Fig. S5a). Next, we compared HRDsum scores between MSI cell lines and microsatellite stable (MSS) cell lines using MSI classifications based on the frequency of deletions within microsatellite regions (Chan et al., 2019). As is the case in patient samples, HRDsum scores were markedly low in MSI cell lines compared to HRDsum scores in MSS cell lines (Fig. 5b). We then performed a correlative analysis to look for associations between HRDsum scores and various genomic features in an unbiased manner. The most significant result was a positive correlation with copy number on chromosome 3q26.2, which peaked at the *ACTRT3* locus (Fig. 5c). *ACTRT3* expression levels also correlated positively with HRDsum scores (Supplementary Fig. S5b). To our knowledge, there is no known biological association between homologous recombination and *ACTRT3* or any of the neighboring hits from the correlative analysis. However, given that telomeric allelic imbalance makes up part of the HRDsum score, it is interesting to note that *ACTRT3* is adjacent to *TERC*, which encodes the RNA component of telomerase. *TERC* was not included in the original copy number analysis, which focused on protein-coding genes. We obtained absolute copy number calls for *TERC* from the DepMap portal (https://depmap.org) and found that, indeed, *TERC* copy number associated with HRDsum scores (Supplementary Fig. S5c). Amplification of 3q26 is characteristic of a class of tumors enriched for *TP53* mutations and copy number changes and was previously shown to associate with HRDsum scores in serous endometrial cancer (Ciriello et al., 2013; Jönsson et al., 2021). Altogether, these observations suggest that key associations with HRDsum scores translate from patient samples to cell lines.

**Figure 5.**
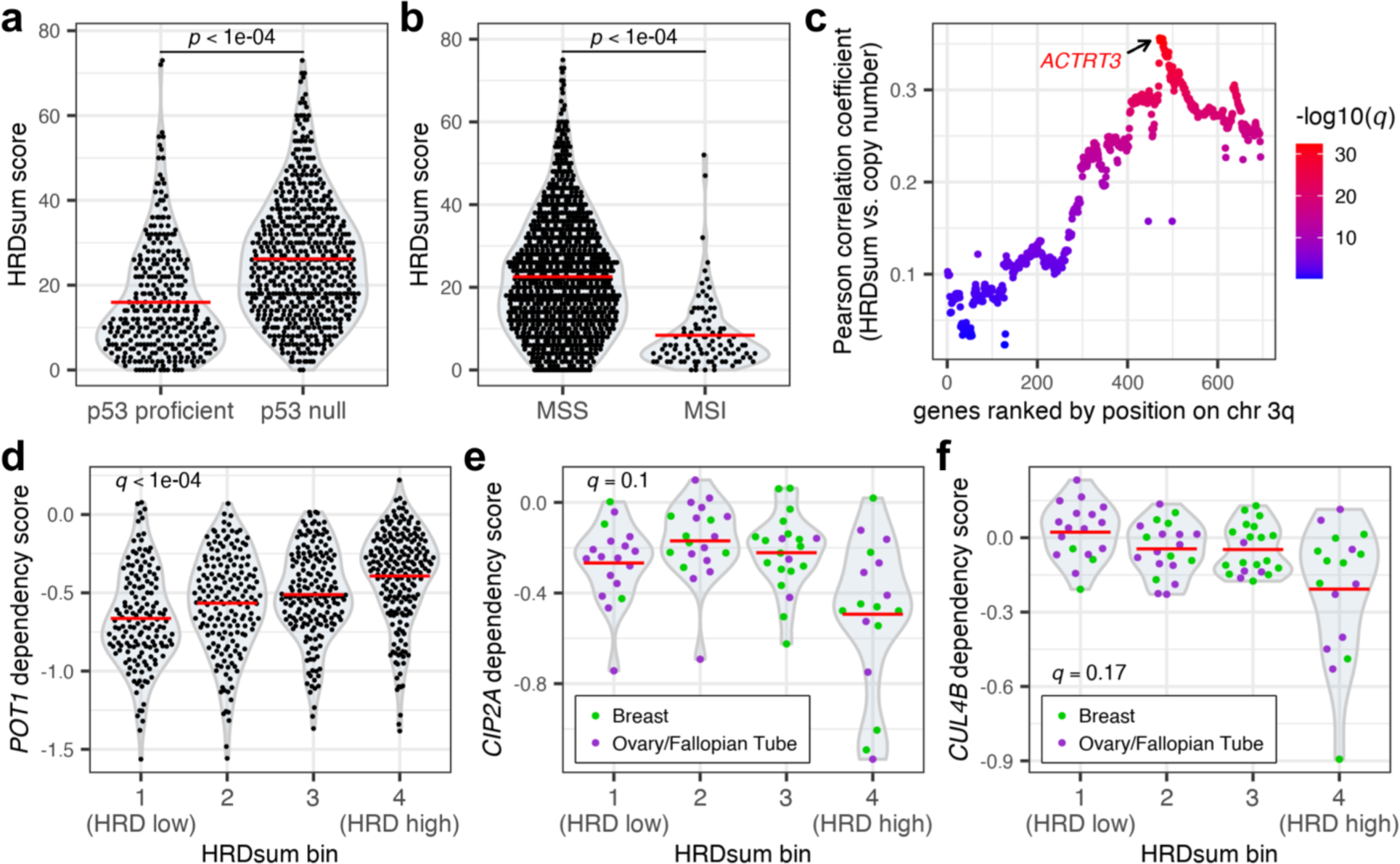
Associations of HRD predictions with genomic features and genetic dependencies. **a)** HRDsum scores in cell lines grouped by p53 functional status. Red bars represent the mean. Mann-Whitney U test *p*-value is shown. **b)** HRDsum scores in cell lines grouped by microsatellite stability classification. Red bars represent the mean. Mann-Whitney U test *p*-value is shown. **c)** Pearson correlation coefficients between HRDsum scores and relative copy number for genes on chromosome 3, arm q (chr 3q). Colors represent Benjamini-Hochberg-adjusted *p*-values (*q*-values) from the Pearson correlation test. **d)** *POT1* dependency scores in cell lines binned by HRDsum score quartiles. Red bars represent the mean. *Q*-value represents the Benjamini-Hochberg-adjusted *p*-value of a Kruskal-Wallis test. **e) – f)** *CIP2A* **(e)** and *CUL4B* **(f)** dependency scores in breast and ovarian cell lines binned by HRDsum score quartiles. Red bars represent the mean. *Q*-values represent the Benjamini-Hochberg-adjusted *p*-values of a Kruskal-Wallis test.

### Associations between HRD classifications and genetic dependencies

To gain new insight into HRD and its therapeutic potential, we took advantage of key datasets that are available exclusively for cell lines. One such dataset is the Cancer Dependency Map (DepMap), which identifies genetic cancer dependencies for over 1,000 cell lines (Pacini et al., 2021; Tsherniak et al., 2017). We used this dataset to determine whether HRD cell lines are particularly vulnerable (or resistant) to certain genetic deficiencies by binning cell lines based on HRDsum scores and then testing for differences in genetic dependency scores. The top two associations were dependency scores for *POT1* and *TINF2*, which encode for two different subunits of the telomere-capping shelterin complex. In particular, cell lines with high HRDsum scores were less sensitive to *POT1* and *TINF2* inactivation than cell lines with low HRDsum scores (Fig. 5d, Supplementary Fig. S5d).

In addition to testing for pan-cancer associations, we performed a similar analysis with only breast and ovarian tissue types, which show the highest frequency of HRD. This approach revealed that dependency on *CIP2A* was most prevalent in the group of cell lines with the highest HRDsum scores (Fig. 5e). The *CIP2A* gene encodes an inhibitor of the protein phosphatase PP2A. *CIP2A* dependency was also associated with CHORD-HRD predictions (Supplementary Fig. S5e). In line with these findings, loss of *CIP2A* was recently reported to be synthetical lethal with BRCA1/2 (Adam et al., 2021). We observed a similar pattern with dependency on *CUL4B*, which encodes part of an E3 ubiquitin ligase complex that participates in the DNA damage response as well as other processes (Y. Li & Wang, 2017; Yi et al., 2015). Sensitivity to *CUL4B* inactivation appeared to be selective, with only a fraction of cell lines showing *CUL4B* dependency, and dependency was specific to the group of cell lines with the highest HRDsum scores (Fig. 5f). Thus, *CUL4B* may harbor potential as a therapeutic target in HRD tumors. More observations are needed to determine whether the association between HRD and *CUL4B* dependency is significant.

### Associations between HRD classifications and drug sensitivities

HRD shows promise as a predictive biomarker for PARP inhibitors (PARPi), as well as platinum-based chemotherapy drugs (Miller et al., 2022; Wiel et al., 2022). To explore this aspect of HRD in cell lines, we compared HRDsum scores to PARPi sensitivity, as determined by a clonogenic assay, for 94 cell lines across 19 different tissue types. Cell lines were categorized as either sensitive or insensitive to PARP inhibition based on cut-offs for both the IC_50_ and maximal activity values (see “Methods”). Overall, HRDsum scores were slightly higher in PARPi-sensitive cell lines than in PARPi-insensitive cell lines (Supplementary Fig. S6a). Looking at specific tissue types, this effect was driven largely by breast cancer lines, which showed a significant difference in HRDsum scores between response groups (Fig. 6a). Ovarian cancer cell lines, on the other hand, showed a wide range of HRDsum scores within the PARPi-sensitive group (Fig. 6a). Outside of breast and ovarian cell lines, there was no difference in HRDsum scores between response groups for cell lines in the remaining 17 tissue types (Fig. 6a). Thus, HRDsum scores were associated with PARPi sensitivity in breast cancer cell lines, but not pan-cancer. We observed a similar, albeit not significant, trend with independent datasets, using either CHORD probabilities for HRD predictions or the PRISM Repurposing dataset for PARPi sensitivity (Supplementary Fig. S6b,c) (Corsello et al., 2020). For platinum-based drugs, there were no associations between HRDsum scores and viability measurements available from the PRISM Repurposing dataset (Fig. 6b, Supplementary Fig. S6d).

**Figure 6.**
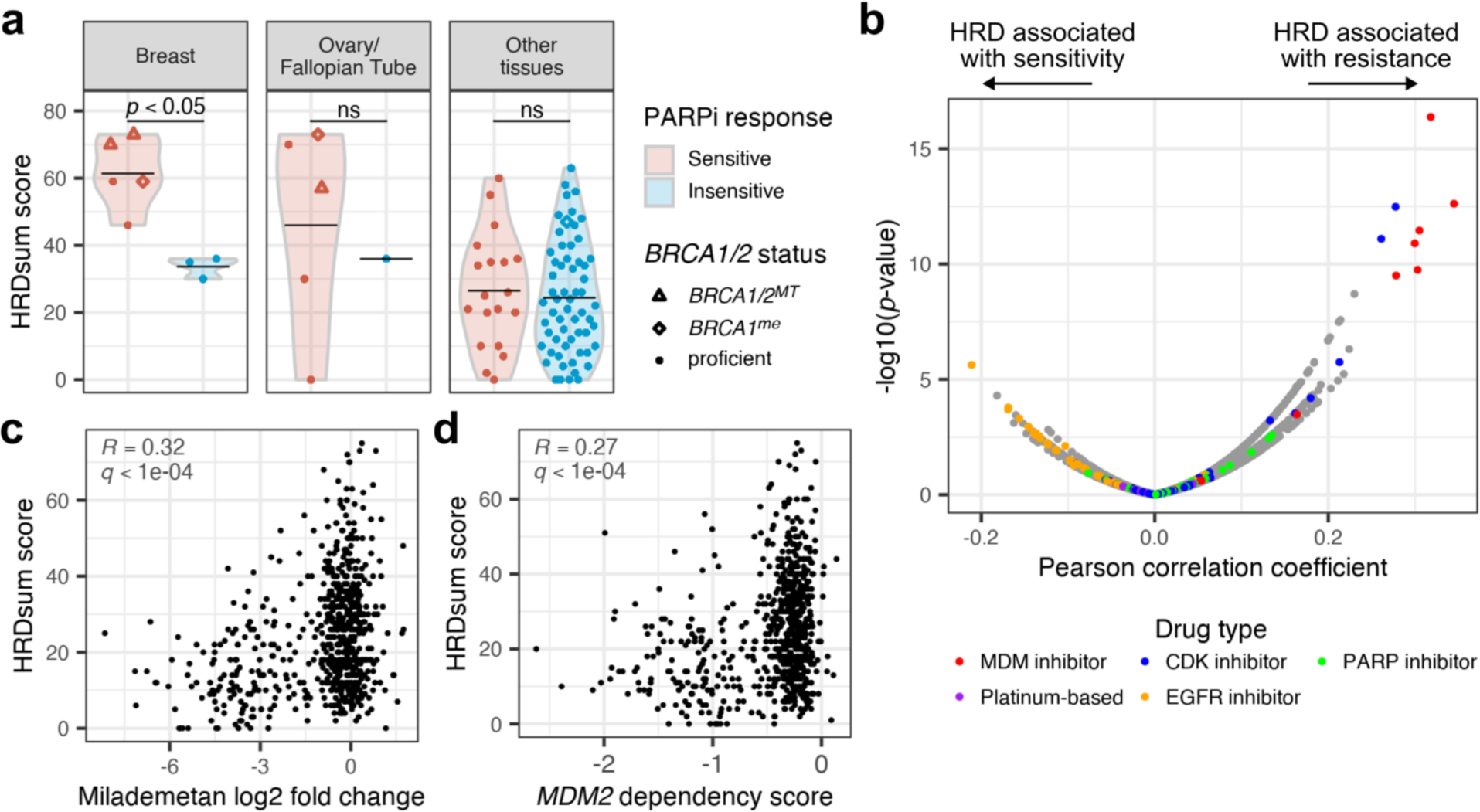
Associations of HRD predictions with drug sensitivities. **a)** HRDsum scores in relation to PARPi response, as determined by a clonogenic assay. Breast and ovarian cancer cell lines are shown in the two left panels, and all other cell lines are shown in the right panel. Point shapes indicate biallelic loss of *BRCA1/2* (*BRCA1/2^MT^*), likely epigenetic silencing of *BRCA1* (*BRCA1^me^*), and likely BRCA1/2 proficiency (proficient). Black bars represent the mean. Mann-Whitney U test *p*-values are shown. ns, not significant. **b)** Pearson correlation coefficients between HRDsum scores and drug sensitivities (log2 fold change in viability for treatment versus DMSO) determined by the PRISM Repurposing 23Q2 dataset. Positive coefficients indicate an association between HRDsum scores and drug resistance, whereas negative coefficients indicate an association between HRDsum scores and drug sensitivity. Colors highlight different drug types of interest. **c)** Correlation between HRDsum scores and sensitivity to the MDM2 inhibitor Milademetan (PRISM Repurposing 23Q2). Sensitivity is shown as the log2 fold change in viability (treatment versus DMSO). Pearson correlation coefficient (*R*) and Benjamini-Hochberg-adjusted *p*-value (*q*) are shown in gray. **d)** Correlation between HRDsum scores and *MDM2* dependency scores (DepMap 22Q4). Pearson correlation coefficient (*R*) and Benjamini-Hochberg-adjusted *p*-value (*q*) are shown in gray.

In addition to testing for previously identified associations, we used the PRISM Repurposing dataset to identify potentially novel associations between HRD predictions and drug sensitivity (Corsello et al., 2020). Interestingly, various MDM2 inhibitors (MDM2i) showed many of the strongest correlations between HRDsum scores and drug response (Fig. 6b). Cell lines sensitive to Milademetan and other MDM2i showed relatively low HRDsum scores, indicating that MDM2i-sensitive lines tend to be HR-proficient (Fig. 6b,c). In line with this observation, cell lines showing a dependency on the *MDM2* gene showed relatively low HRDsum scores (Fig. 6d). MDM2 negatively regulates p53, and the primary mechanism of action for MDM2i involves activation of the p53 pathway (Vassilev et al., 2004). Given that response to MDM2i depends on functional p53 and that HRD cell lines are typically p53-deficient, we reasoned that *TP53* loss underlies the resistance of HRD cell lines to MDM2i. Consistent with this idea, HRD was mutually exclusive with dependence on *MDM4* and *PPM1D*, which, like *MDM2*, also negatively regulate p53 (Supplementary Fig. S7a) (Wasylishen & Lozano, 2016). In addition to MDM2i, HRDsum scores were also associated with resistance to CDK inhibitors, particularly those targeting CDK4 and CDK6 (CDK4/6i) (Fig. 6b, Supplementary Fig. S7b). This association may also be explained by p53 loss in HRD cell lines, as there is evidence to suggest that response to CDK4/6 inhibition is p53-dependent (Bellutti et al., 2018; Wang et al., 2022). On the other end of the spectrum, significant negative correlates were predominantly EGFR/HER2 inhibitors (Fig. 6b). However, these correlations were relatively weak, and cell lines with pathogenic *EGFR* mutations did not have particularly high HRDsum scores, which one would expect if HRD were predictive of sensitivity to EGFR inhibition (Supplementary Fig. S7c). In conclusion, a survey of PRISM drug responses revealed that HRD associated with resistance to MDM2i and, to a lesser extent, CDK4/6i, and did not associate well with sensitivity to any drugs. Similar trends were observed with drug responses available from the GDSC2 dataset (Supplementary Fig. S7d) (Yang et al., 2013).

## DISCUSSION

Multiple methods based on mutational scars have been developed and used to predict HRD for large cohorts of patient samples. However, equivalent datasets for cell lines have been lacking. In this study, we determined HRD predictions for over 1,400 cell lines using both a traditional method based on HRDsum scores, as well as the more recently developed classifier CHORD. Given that HRDsum scores and CHORD were designed to analyze mutational patterns in patient tumor samples, it was initially unclear how well these methods would perform on cell lines for multiple reasons. First, the mutational profiles of cell lines can differ from tumors, as cell lines are less heterogeneous and tend to harbor more mutations overall (Goodspeed et al., 2016). Second, HRDsum scores and CHORD rely specifically on somatic mutations, which are less straightforward to detect in cell lines since most cancer cell lines lack matching normal cell lines. For HRDsum scores, which can be determined from SNP arrays or WES, somatic mutations can be inferred by using a pool of non-matching normal cell lines a reference. CHORD, however, requires WGS, which is not readily available for many normal cell lines. To circumvent this issue, we applied a series of filters designed to remove common variants that are most likely germline or artifacts of PCR amplification or sequencing. Despite the aforementioned differences between cell lines and patient samples, HRDsum scores and CHORD were able to discriminate BRCA1/2-deficient cell lines from BRCA1/2-proficient cell lines. Furthermore, HRD predictions in cell lines showed several similarities to HRD in patient samples, including an association with p53 loss and enrichment in breast and ovarian cancer types. Overall, these results suggest that genomic scar-based methods may be utilized for predicting HRD in cell lines and that cancer cell lines are representative of tumors with respect to key HRD characteristics. As WGS becomes more available, this adaptation of CHORD for cell lines can be readily applied to additional samples.

HRDsum scores and CHORD probabilities agreed well overall, particularly for BRCA1/2-deficient cell lines. Some cell lines, however, were classified as HRD in only one of the two datasets. One possible explanation for the differences observed between the two datasets is that CHORD and HRDsum scores each capture different nuances in the DNA repair landscape, which can vary depending on the genetic or epigenetic background of a tumor or cell line. The following points support this idea. First, CHORD and HRDsum scores detect different types of mutations: the top predictors for the CHORD model are small deletions with flanking microhomology and duplications, whereas HRDsum scores are based on large scale changes in allelic content and copy number. Second, given that the loss of HR factors can result in different types of DNA damage that can be repaired by a variety of pathways, different cell lines may acquire different types of mutational scars depending on the nature of the HR deficiency and/or the functional status of compensatory repair pathways (Koh et al., 2021). Thus, HRD is a complex phenotype in terms of mutational consequences, and different HRD detection methods are likely to detect different subtleties of this phenotype.

Compared to other HRD detection methods, genomic scar-based methods such as CHORD and HRDsum scores have many advantages as well as limitations. Since HRD-induced scars are a relatively permanent phenotype that will persist even if HR is restored, these scars may not always represent the current status of HR. This was indeed the case for the *BRCA2*-revertant cell line HCC1428, which was classified as HRD by both CHORD and the HRDsum method. However, since genomic scars are a downstream consequence of HRD, scar-based methods have a major advantage of being able to detect etiologies of HRD that extend beyond the loss of BRCA1 and BRCA2. Interestingly, the two FANCC-deficient cell lines included in our study were classified as HRD by either CHORD or HRDsum scores. FANCC is part of the FA core complex, which has been shown to promote HR (Nakanishi et al., 2005). Therefore, FANCC loss potentially explains the HRD phenotype of a subset of cell lines. Additional *FANCC*-mutant cell lines will need to be assessed to determine whether FANCC loss significantly associates with HRD. Probable explanations for HRD remain undetermined for some cell lines such as CAOV3, which scored high for HRD in both the CHORD and HRDsum datasets, but does not show biallelic loss of any DNA damage repair-related genes other than *TP53*. It is possible that certain monoallelic mutations or epigenetic alterations, either alone or in combination, underlie HRD in such cell lines.

In addition to validating previously observed associations between HRD predictions and various features, cell line-specific datasets such as the DepMap dependency scores enabled us to gain new insights into HRD. We found, for instance, that HRD negatively associated with genetic dependencies on POT1 and TINF2, two subunits of the shelterin complex. POT1 was recently shown to preserve the stability of telomeres by protecting them from unwanted HR activity (Glousker et al., 2020). Thus, POT1 inactivation is more detrimental when HR is intact, which could explain why HR-proficient cell lines were more sensitive to loss of POT1. We also found that *CUL4B* dependency was specific to HRD cell lines in breast and ovarian cancer types. Loss of CUL4B sensitizes cells to DNA damaging agents and is thought to accelerate apoptosis in response to DNA damage (Yi et al., 2015). It would therefore be interesting to further explore whether CUL4B loss is synthetic lethal with HRD. Given that CUL4B is amenable to small molecule inhibition and appears to be selectively essential in HRD cell lines, CUL4B could serve as an attractive therapeutic target.

HRD is actively being explored as a predictive biomarker for sensitivity to PARP inhibition, and two different genomic scar-based HRD assays, Myriad MyChoice CDx and Foundation Medicine %LOH, have been approved as diagnostics for PARPi therapy (Ngoi & Tan, 2021). In clinical studies including PAOLA-1, ARIEL2, and ARIEL3, HRD patient groups showed a greater benefit from PARPi treatment than HR-proficient patient groups (Coleman et al., 2017; Ray-Coquard et al., 2019; Swisher et al., 2017). However, current HRD assays have yet to consistently discriminate responders from non-responders, suggesting that HRD-associated genomic scars are an imperfect biomarker of PARPi sensitivity (Chiang et al., 2021; Ladan et al., 2021). In cell lines, we found that HRD predictions associated with PARPi sensitivity in breast cancer, but not other cancer types. Lack of a pan-cancer association between HRD and PARPi sensitivity was also reported in a recent study of CCLE cell lines (Takamatsu et al., 2024). More work needs to be done to fully understand the relationship between HRD and PARPi response, and we hope that the resources provided in this study facilitate this effort.

We envision a wide range of future applications for the datasets generated by this work. In this study, we performed univariate association analyses to provide an initial characterization of HRD in cell lines. However, multivariate analyses could yield additional insights and tools. For instance, the HRDsum scores could be used to build a gene expression-based classifier of HRD, which would enable the prediction of HRD in samples where only RNA sequencing is available. We also anticipate that the HRD predictions generated in this study will aid in the selection of cell lines for experiments relating to DNA repair. Potential applications of this work extend beyond the study of HRD, as well, as we also provide a table of WGS-based mutation contexts, or counts of different mutation types, for over 300 cell lines. Only a subset of these mutation types were utilized by CHORD to predict HRD, and it may be worth exploring other mutation types or combinations thereof to determine their potential to predict other types of DNA repair deficiencies, for example. In conclusion, further exploration of the datasets from this study is warranted and should advance our understanding of DNA repair deficiencies in cancer.

## ACKNOWLEDGMENTS

All studies were funded by KSQ Therapeutics.

