## Supplementary Figures for "Pan-cancer Analysis of Homologous Recombination Deficiency in Cell Lines"

#### Supplementary Figure S1

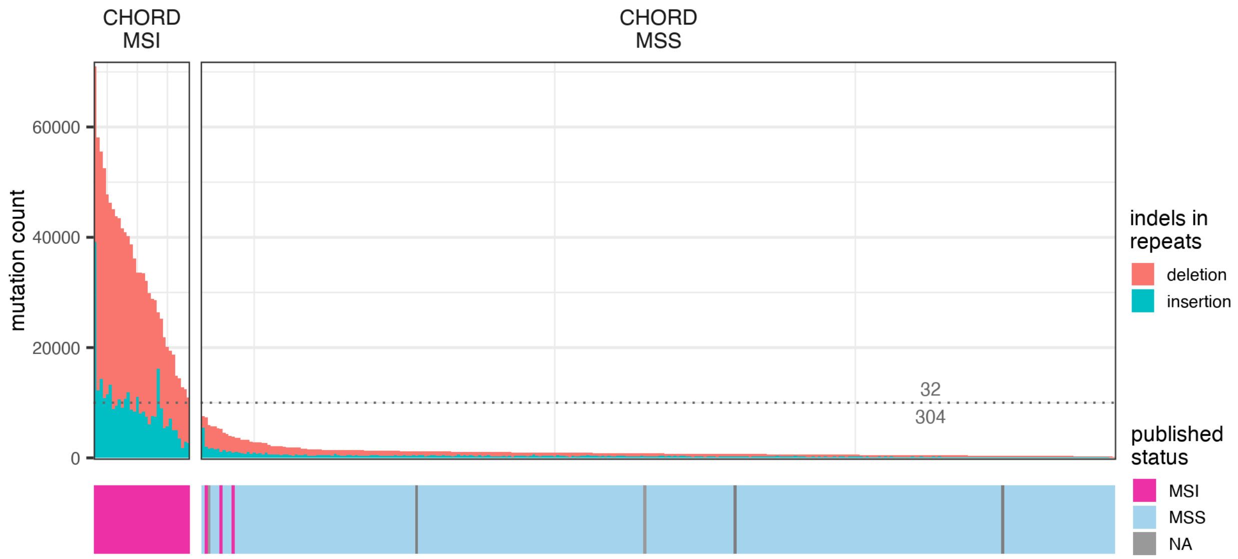


**Supplementary Figure S1. CHORD MSI predictions.** Top plot: Cell lines ranked by the number of indels present in repetitive regions. Cell lines with > 10,000 indels in repetitive regions were classified by CHORD as showing microsatellite instability (CHORD MSI) and were therefore ineligible for HRD assessment by CHORD. The other 304 cell lines were classified by CHORD as showing microsatellite stability (CHORD MSS). Bottom plot: MSI and MSS calls determined in a previous study (Chan et al., 2019). NA, MSI status not available.

### Supplementary Figure S2

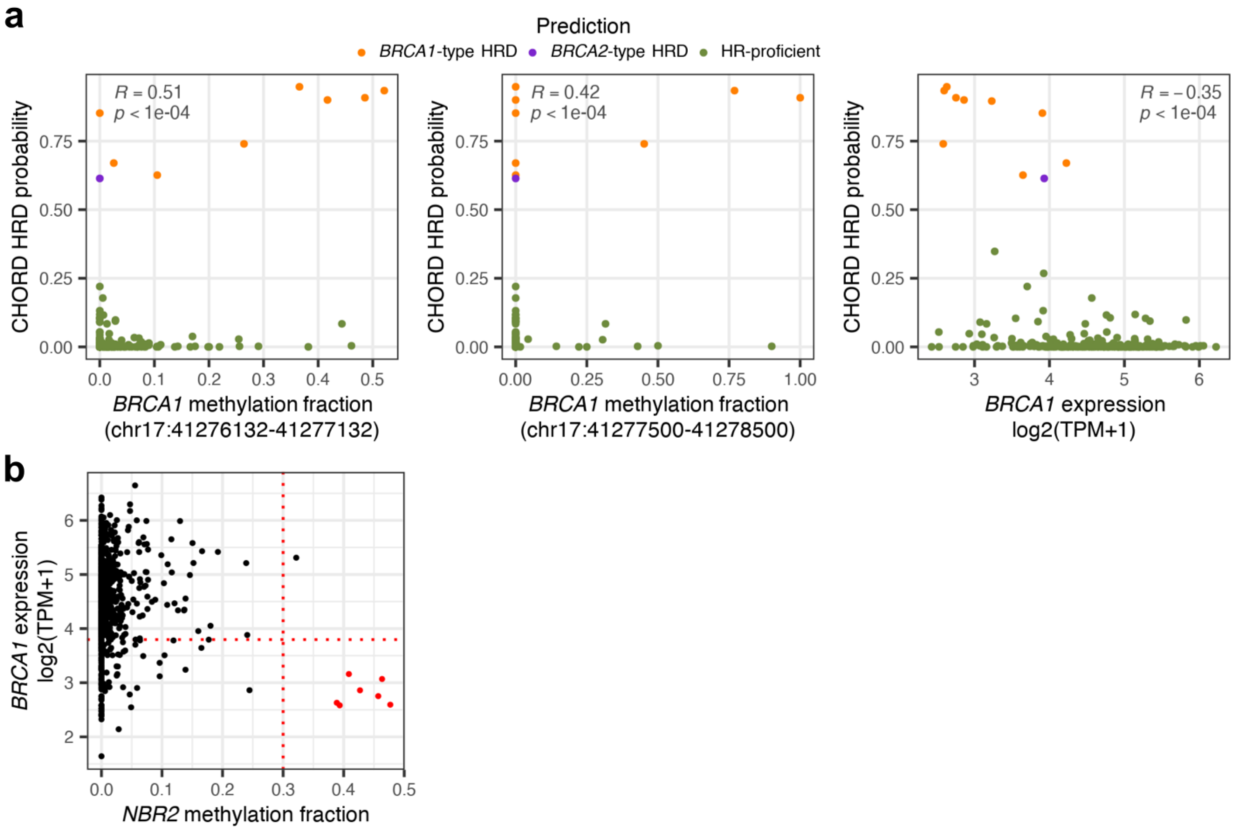


**Supplementary Figure S2. CHORD predictions relative to *BRCA1* expression state. a)** Correlations of CHORD HRD probabilities with *BRCA1* promoter methylation (left and middle panels) and *BRCA1* gene expression (right panel). Colors represent CHORD classifications. Pearson correlation coefficient (*R*) and Benjamini-Hochberg-adjusted *p*-values are shown in gray. **b)** Identification of cell lines showing evidence of *BRCA1* gene silencing. Likely cases of epigenetic silencing of *BRCA1* were determined by selecting cell lines with *NBR2* promoter methylation fraction > 0.3 and *BRCA1* expression levels below the lower quartile (red points).

### Supplementary Figure S3

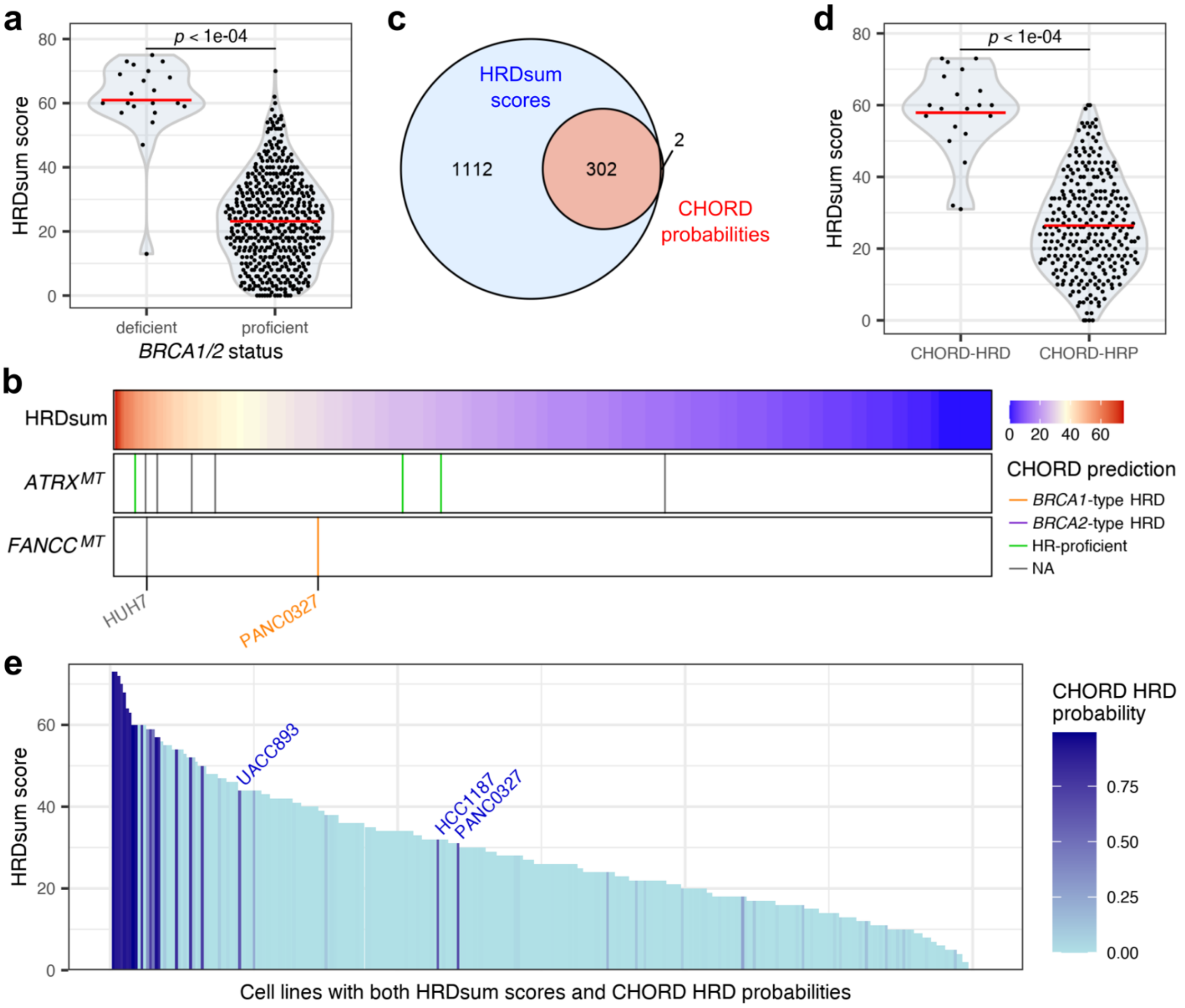


**Supplementary Figure S3. Characterization of HRDsum scores in cell lines. a)** HRDsum scores in cell lines grouped by BRCA1/2 functional status. BRCA1/2 deficiency was defined as either biallelic loss of *BRCA1/2* or likely epigenetic silencing of *BRCA1*. Red bars represent the mean. Mann-Whitney U test *p*-value is shown. **b)** *ATRX* and *FANCC* mutations in cell lines ranked by HRDsum score. Only biallelic loss-of-function mutations are shown. Mutant cell lines are colored by CHORD predictions, if available. NA, not available. **c)** Venn diagram showing the overlap in cell lines between the CHORD and HRDsum datasets. **d)** HRDsum scores in cell lines grouped by CHORD prediction. Red bars represent the mean. Mann-Whitney U test *p*-value is shown. **e)** Comparison of HRDsum scores (Y-axis) and CHORD HRD probabilities (color) for cell lines present in both datasets.

### Supplementary Figure S4

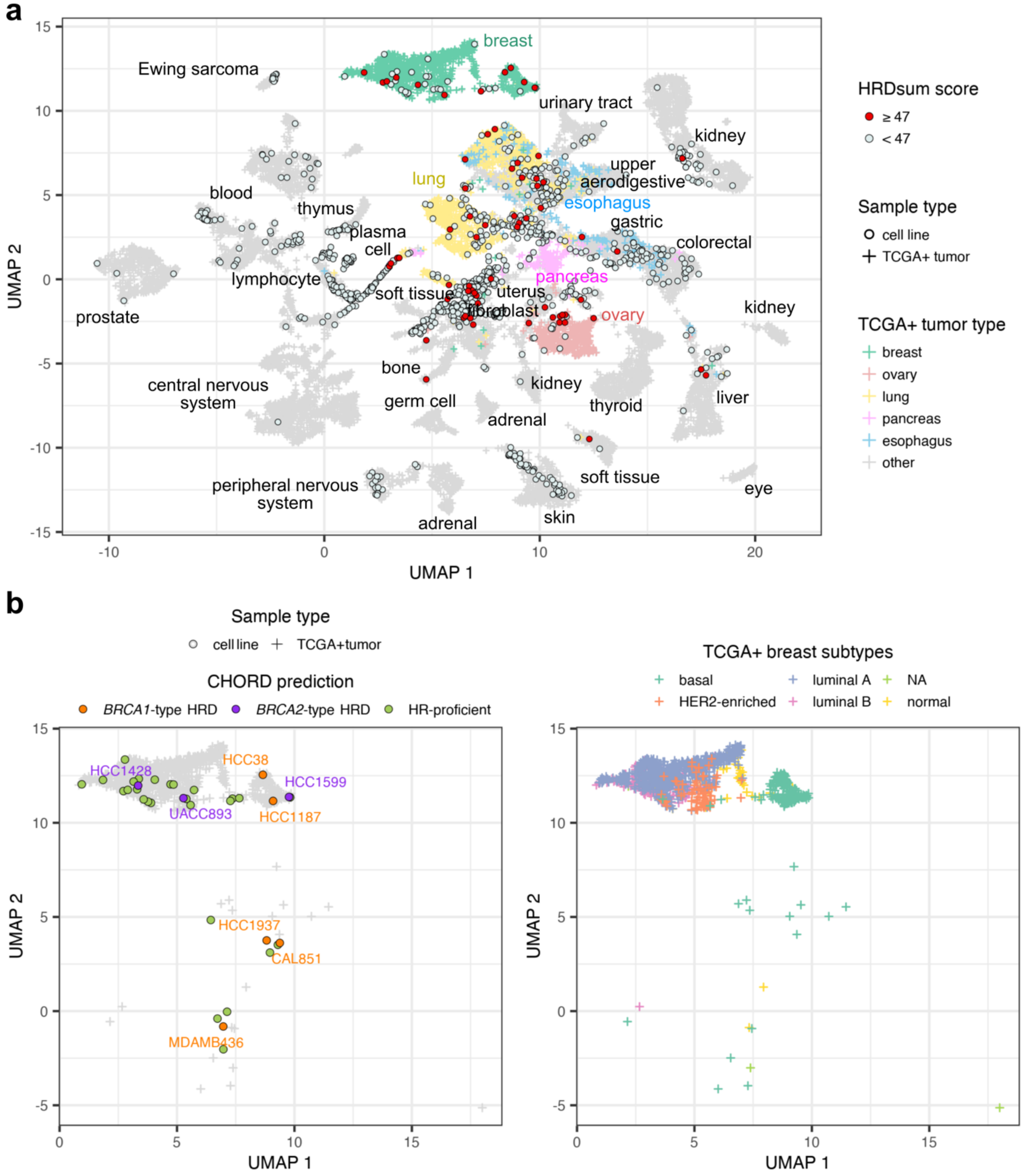


**Supplementary Figure S4. Alignment of HRD predictions to tumor types using Celligner. a)** Uniform Manifold Approximation and Projection (UMAP) plot of the Celligner dataset colored by HRDsum predictions. **b)** Left panel: UMAP plot of Celligner dataset for breast cancer samples colored by CHORD predictions. Right panel: UMAP plot of Celligner dataset for TCGA+ breast cancer samples colored by subtype.

### Supplementary Figure S5

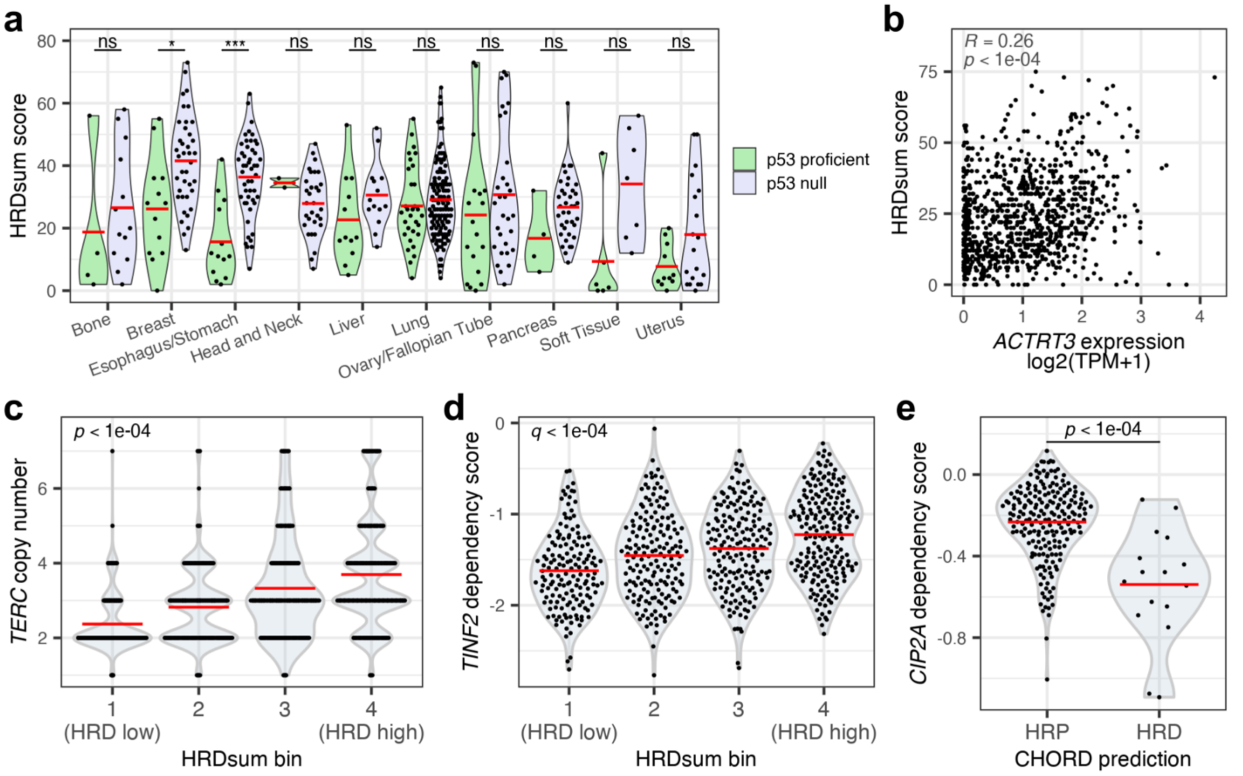


**Supplementary Figure S5. Additional associations of HRD predictions with genomic features and genetic dependencies. a)** HRDsum scores in p53-proficient cell lines versus p53-null cell lines, grouped by tissue type. Only tissue types with at least one HRDsum-high cell line are shown. Red bars represent the mean. ***, Benjamini-Hochberg-adjusted *p*-value < 0.001; *, Benjamini-Hochberg-adjusted *p*-value < 0.05; ns, not significant (Mann-Whitney U test). **b)** Correlation between HRDsum scores and *ACTRT3* expression (DepMap 22Q4). Pearson correlation coefficient (*R*) and *p*-value are shown in gray. **c)** *TERC* absolute copy number in cell lines binned by HRDsum score quartiles. Red bars represent the mean. Kruskal-Wallis test *p*-value is shown. **d)** *TINF2* dependency scores in cell lines binned by HRDsum score quartiles. Red bars represent the mean. *Q*-value represents the Benjamini-Hochberg-adjusted *p*-value of a Kruskal-Wallis test. **e)** *CIP2A* dependency scores in CHORD-HRP cell lines versus CHORD-HRD cell lines. Red bars represent the mean. Mann-Whitney U test *p*-value is shown.

### Supplementary Figure S6

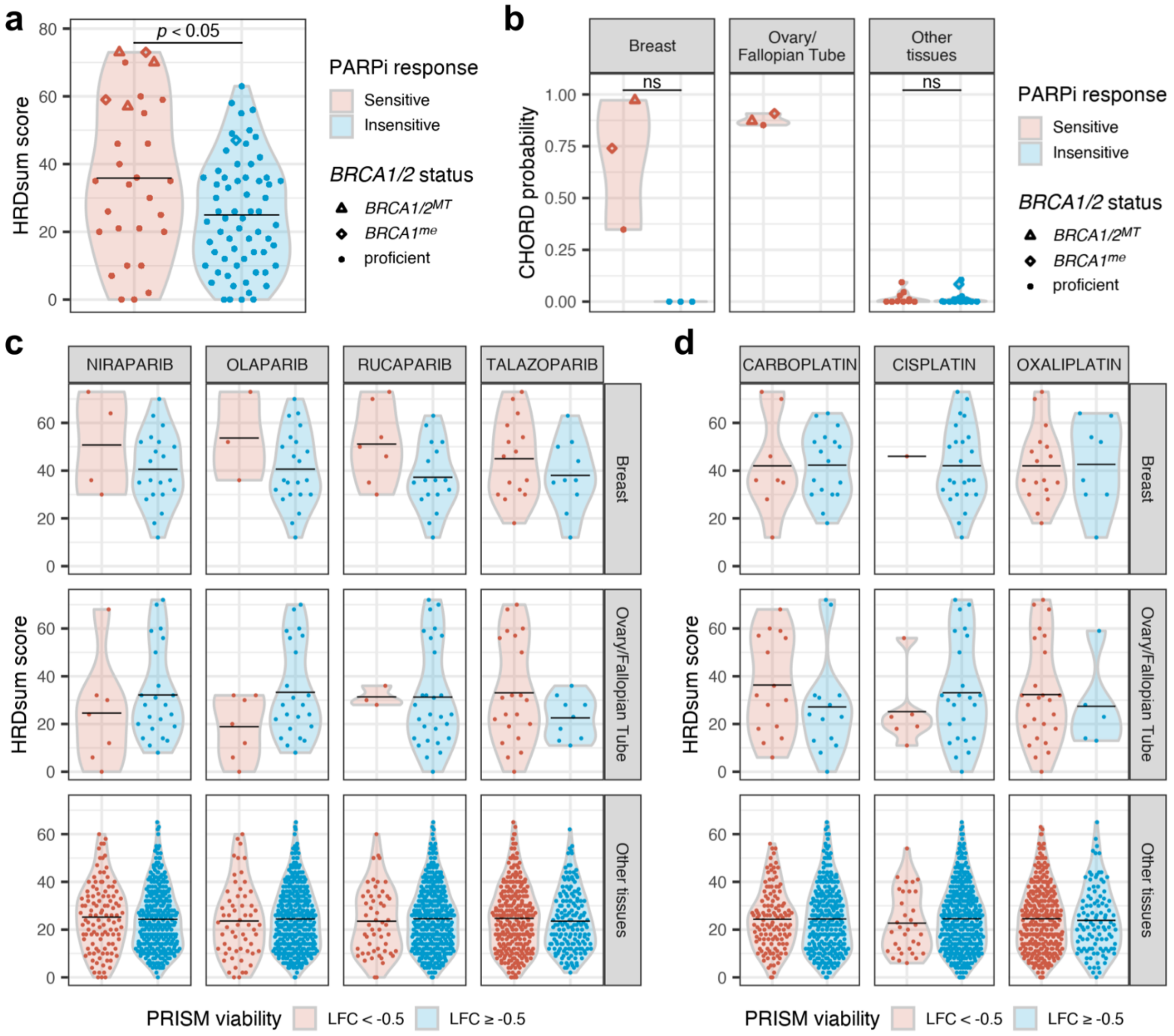


**Supplementary Figure S6. HRD predictions in relation to PARP inhibitor and platinum-based treatment response. a) – b)** HRD predictions in cell lines grouped by response to PARP inhibition, as measured by a clonogenic assay. Point shapes indicate biallelic loss of *BRCA1/2* (*BRCA1/2*$^{MT}$), likely epigenetic silencing of *BRCA1* (*BRCA1*$^{me}$), and likely BRCA1/2 proficiency (proficient). Black bars represent the mean. Mann-Whitney U test *p*-values are shown. ns, not significant. **c) – d)** HRDsum scores in cell lines grouped by response to various PARP inhibitors **(c)** and platinum-based chemotherapy drugs **(d)** using the PRISM Repurposing dataset (23Q2). Cell lines were grouped based on the log2 fold change in viability (treatment versus DMSO) using a cut-off of -0.5. Black bars represent the mean. None of the comparisons were significant (Mann-Whitney U test).

### Supplementary Figure S7

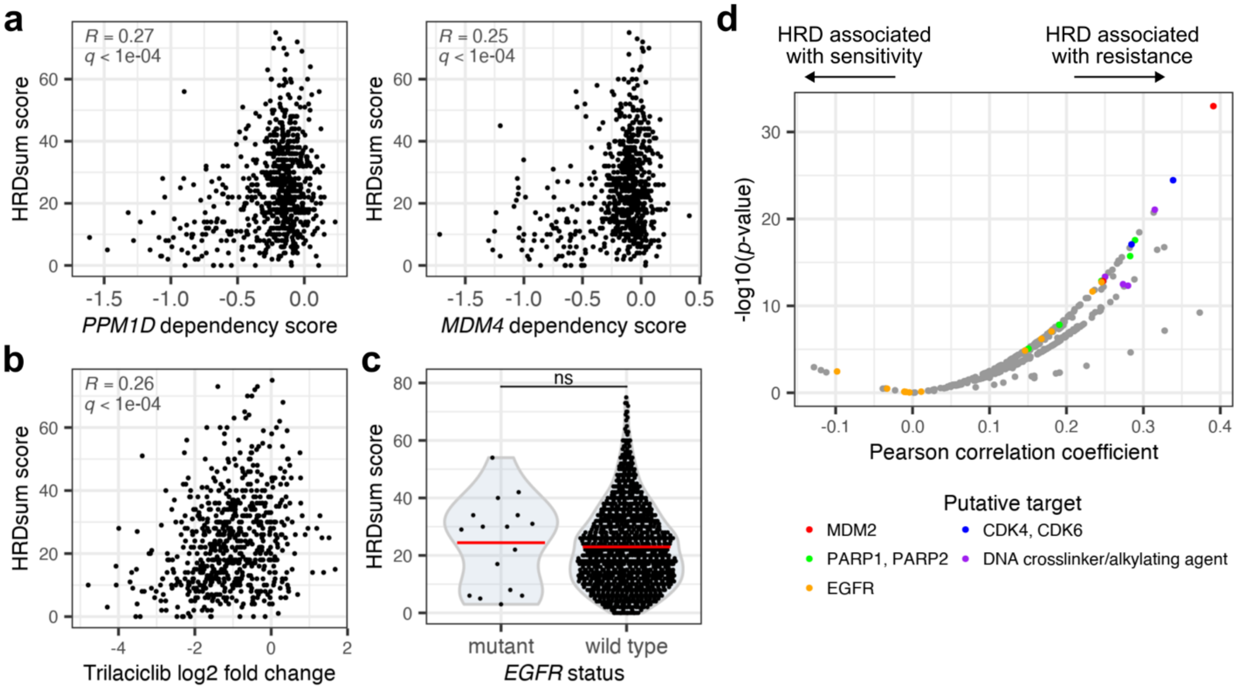


**Supplementary Figure S7. Additional analyses related to associations of HRD predictions with drug sensitivities. a)** Correlations between HRDsum scores and either *PPM1D* or *MDM4* dependency scores (DepMap 22Q4). Pearson correlation coefficients (*R*) and Benjamini-Hochberg-adjusted *p*-values (*q*) are shown in gray. **b)** Correlation between HRDsum scores and sensitivity to the CDK4/6 inhibitor Trilaciclib (PRISM Repurposing 23Q2). Sensitivity is shown as the log2 fold change in viability (treatment versus DMSO). Pearson correlation coefficient (*R*) and Benjamini-Hochberg-adjusted *p*-value (*q*) are shown in gray. **c)** HRDsum scores in relation to *EGFR* genotype. Cell lines harboring an *EGFR* mutation annotated as "pathogenic", 'likely pathogenic", and/or "drug response" in ClinVar were classified as "mutant". Cell lines lacking detectable variants in *EGFR* were classified as "wild type". Red bars represent the mean. ns, not significant (Mann-Whitney U test). **d)** Pearson correlation coefficients between HRDsum scores and the log10($IC_{50}$) for drugs available from the GDSC2 dataset. Positive coefficients indicate an association between HRDsum scores and drug resistance, whereas negative coefficients indicate an association between HRDsum scores and drug sensitivity. Colors highlight different drug targets of interest.

### SUPPLEMENTARY TABLES

**Supplementary Table S1. Mutation contexts for cell lines with WGS available.** Mutation contexts were extracted for SNVs, indels, and structural variants using the CHORD package in R (L. N. Nguyen, 2022).

**Supplementary Table S2. CHORD predictions for cell lines.** Includes the output of CHORD as well as cell line metadata. The "hr_status" and "hrd_type" columns were adjusted to an HRD cut-off of 0.4.

**Supplementary Table S3. Confusion matrix of CHORD HRD predictions in cell lines.** Labeled and predicted HRD classes using a CHORD cut-off of 0.4.

**Supplementary Table S4. DNA repair mutation enrichment analysis for CHORD predictions.** The "pct_hrd" and "pct_hrp" columns indicate the percent of CHORD-HRD and CHORD-HRP cell lines showing a deficiency in each gene, respectively. *P*-values and the Benjamini-Hochberg-adjusted *p*-values (*q*-values) are from a one-tailed Fisher's Exact test. Only DNA repair genes with a deficiency in at least one CHORD-HRD cell line were analyzed.

**Supplementary Table S5. HRDsum scores for cell lines.** Columns beginning with "hrdsum" contain the raw HRDsum scores for the "Broad", "Sanger (Sanger WES)", and "Sanger (Broad WES)" datasets and the summary HRDsum scores derived from merging all three datasets.

**Supplementary Table S6. DNA repair mutation enrichment analysis for HRDsum scores.** The "pct_hrd" and "pct_hrp" columns indicate the percent of HRDsum-high and non-HRDsum-high cell lines showing a deficiency in each gene, respectively. *P*-values and the Benjamini-Hochberg-adjusted *p*-values (*q*-values) are from a one-tailed Fisher's Exact test. Only DNA repair genes with a deficiency in at least one HRDsum-high cell line were analyzed.
